# Polyadenylation of mRNAs encoding secreted proteins by TENT5 family of enzymes is essential for gametogenesis in mice

**DOI:** 10.1101/2023.08.01.551432

**Authors:** Michał Brouze, Agnieszka Czarnocka-Cieciura, Olga Gewartowska, Monika Kusio-Kobiałka, Kamil Jachacy, Marcin Szpila, Bartosz Tarkowski, Jakub Gruchota, Paweł Krawczyk, Seweryn Mroczek, Ewa Borsuk, Andrzej Dziembowski

## Abstract

Cytoplasmic polyadenylation plays a vital role in gametogenesis, however, the participating enzymes and substrates in mammals remain unclear. Using knockout and knock-in mouse models, we describe the essential role of 4 TENT5 poly(A) polymerases in mice fertility and gametogenesis.

TENT5B and TENT5C play crucial, but redundant roles in oogenesis, with double knockout of both genes leading to oocyte degeneration. Additionally, TENT5B-GFP knock-in females display gain-of-function infertility effect with multiple chromosomal aberrations in ovulated oocytes. TENT5C and TENT5D both regulate different stages of spermatogenesis, shown by sterility of males with either gene’s knockout mutation. Finally, *Tent5a* knockout significantly lowers fertility, although the underlying mechanism is not directly related to gametogenesis.

Through Direct RNA sequencing we discovered that TENT5s polyadenylate mRNAs encoding endoplasmic reticulum-targeted proteins essential for gametogenesis. Sequence motives analysis and reporter mRNA assay revealed that the presence of endoplasmic reticulum-leader represents the primary determinant of TENT5-mediated regulation.

## Introduction

Essentially every eukaryotic mRNA possesses poly(A) tail which, bound by poly(A) binding proteins (PABPs), facilitates mRNA export from the nucleus and enhances protein synthesis through interactions with translation initiation factors. Poly(A) tail also stabilizes mRNA molecules by preventing exoribonucleolytic decay. Consequently, the deadenylation rate largely determines mRNA half-life. When a poly(A) tail in the cytoplasm is shortened to ∼20 nt, PABP is released, and the mRNA becomes translationally inactive and susceptible to degradation. However, in specific contexts, deadenylated mRNA can be stored in a dormant state to be later re-adenylated in the cytoplasm to activate protein synthesis^1-4^. Such non-canonical cytoplasmic polyadenylation was mostly studied in the context of gametogenesis^1-4^ and local translation at synapses^5,6^. In these instances, certain mRNAs are rapidly polyadenylated in response to cellular signals, which allows translation to start in a transcription-independent fashion. Our knowledge about cytoplasmic polyadenylation comes mostly from either biochemical analyses in *Xenopus laevis* oocytes or genetic studies in invertebrates. The first and only extensively studied cytoplasmic poly(A) polymerase is GLD2 (TENT2 according to the new nomenclature)^3,5,7-14^. In non-mammalian species, including *Caenorhabditis elegans*, *Xenopus laevis* and *Drosophila melanogaster,* TENT2 clearly participates in the elongation of poly(A) tails in gametes and early embryos^3,4,10,15^. TENT2 is also important for synaptic plasticity in fruit fly^5^. On its own TENT2 does not contain any detectable RNA binding domain and in order to select substrates, it cooperates with several RNA binding proteins, like *C. elegans’* GLD3^4^. Seminal work on *X. laevis* oocytes led to a model of cytoplasmic polyadenylation where cycles of deadenylation and TENT2-mediated polyadenylation are regulated by the cytoplasmic polyadenylation element binding protein 1 (CPEB1), in cooperation with several other factors^16^. Based on the data, it seemed very likely that TENT2 is involved in cytoplasmic polyadenylation in mammals. Further support came from experiments where human TENT2 tethered to a reporter mRNA and injected into *X. laevis* oocytes activated translation. In line with this hypothesis knock-down or overexpression of TENT2 in mice oocytes results in delayed maturation and frequent arrest in metaphase I^17^. However, surprisingly, TENT2 knockout (KO) mice of both sexes are fertile and display no major phenotype^18^. Oocyte maturation is normal and poly(A) tail length of reporter mRNA is not altered, neither in germ line nor in somatic cells^18^. Moreover, TENT2 KO mice do not exhibit any behavioral abnormalities, suggesting that polyadenylation in neurons is also unaffected^7^.

This raises a possibility that in mammals, other TENT protein(s), as yet unidentified, could be involved in poly(A) length regulation. However, all other previously described mammalian non-canonical poly(A) polymerases are either mitochondrial (mtPAP)^19^ or mostly nuclear (TENT4A and TENT4B)^19-22^, unlikely to play a significant role in cytoplasmic polyadenylation. In contrast to TENT2, its potential mammalian regulators, CPEBs (1 to 4), were more comprehensively studied^23^. Indeed, CPEB1 KO mice display severe gametogenesis defects^24^ as well as impaired long-term potentiation in neurons^25^. Moreover, other three CPEB proteins contribute to regulation of various physiological processes, although in many cases, they function as translational repressors rather than activators^26^. In conclusion, the mechanisms and the impact of cytoplasmic polyadenylation during gametogenesis remain to be established.

A few years ago we described a previously overlooked family of non-canonical poly(A) polymerases: TENT5 (FAM46)^27^, all members of which were predicted to contain a putative nucleotidyltransferase catalytic domain^27,28^. In mammals, this family of cytoplasmic proteins has four members (TENT5A, TENT5B, TENT5C and TENT5D)^28^ that are differentially expressed in tissues and organs. TENT5A is involved in osteogenesis^29^, TENT5C enhances expression of immunoglobulin mRNA in B cells^30^, while TENT5A and TENT5C together regulate the expression of innate immune response proteins in macrophages^31^. Here, we analyzed gametogenesis-related phenotypes of all four *Tent5* KO mutations. We demonstrate the involvement of TENT5B and TENT5C in oogenesis. We further show that TENT5C and TENT5D participate in spermatogenesis, which was also recently identified by others^32,33^. Notably, using Direct RNA sequencing, we identified TENT5 substrates in both ovaries and testes. The most prominent mRNAs regulated by TENT5 encode secreted proteins, many of which are essential for gametogenesis like zona pellucida components produced by oocytes. The leader sequence for endoplasmic reticulum (ER) is also sufficient for TENT5-mediated poly(A) tail length regulation in oocytes, pointing to a simple mechanism of substrate recognition.

## Results

### TENT5 poly(A) polymerases are involved in both spermatogenesis and oogenesis

To understand the physiological role of the TENT5 family of cytoplasmic poly(A) polymerases we generated and analyzed genetically modified mouse lines carrying constitutive knockout mutations in all *Tent5* family members. The *Tent5a* and *Tent5c* KO mouse lines were describedpreviously^27,29^The *Tent5b* KO mouse line has an 11bp deletion leading to a p.S119EfsX15 frameshift mutation. Simultaneously, we developed another *Tent5b* KO mouse line with a deletion spanning the complete catalytic center of the protein (p.L121P, G122_S238del). Both mouse lines exhibited an identical phenotype, hence they were used interchangeably in this study. The *Tent5d* KO mouse line has a del36bp, ins6bp mutation resulting in the deletion of 10 amino acids, including D82 and D84, which are critical for TENT5D activity, and the insertion of two amino acids (p.76-85del, insCL). Furthermore, due to the lack of reliable, commercially available specific antibodies against TENT5 proteins, we generated several new knock-in mouse lines with either endogenous TENT5B or TENT5D tagged with GFP or FLAG tags. Also, we used the previously generated FLAG- and GFP-tagged TENT5C mouse lines^27,30^.

Individual KOs of all *Tent5* genes did not impact the development and birth ratio of offspring. As *Tent5d* is X chromosome-located, homozygotic KO females could not be generated due to male infertility described further (Figure 1. A, *Tent5a* data previously reported in Gewartowska *et al.* ^29^). Moreover, except previously described *Tent5a* KO line^29^, they also did not affect overall weight (Supplemental Figure 1. A) or except *Tent5c* KO^27^, blood morphology (Supplemental Figure 1. B).

**Figure 1.**
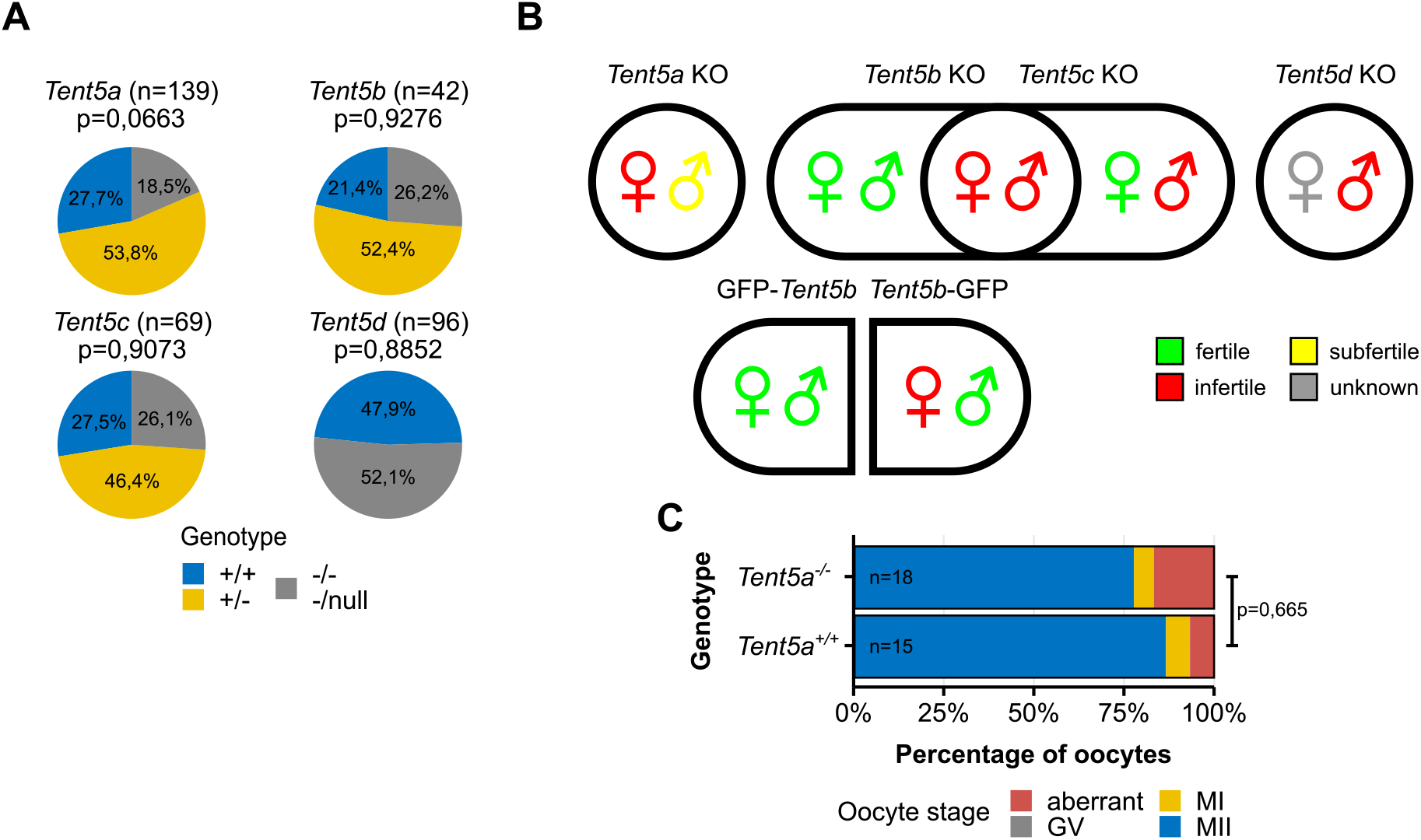
TENT5 poly(A) polymerases are involved in both spermatogenesis and oogenesis. **A.** Pups genotype distribution in all *Tent5* KO mouse lines litters (only males were accounted for in *Tent5d* KO line). P-values reported for comparison with expected Mendelian ratios in Fisher exact test, two-tailed; n values represent a number of pups genotyped. **B.** Fertility status of homozygotic males and females of all studied *Tent5* KO and *Tent5b* GFP-tagged mouse lines. **C.** *In vitro* maturation efficiency of *Tent5a^+/+^*and *Tent5a^-/-^* GV oocytes. P-value reported for comparison of the ratio of metaphase II oocytes to other oocytes after 24h of *in vitro* culture in Fisher exact test, two-tailed; n values represent the number of oocytes analyzed; GV = germinal vesicle, MI = metaphase I, MII = metaphase II

During the initial breeding process, while establishing KO mouse lines, we observed that homozygotic KO mutations of several *Tent5* genes caused partial or complete infertility of both males and females (Figure 1. B): complete male, but not female, infertility of *Tent5c* KO (*Tent5c^-/-^)*; male fertility of *Tent5d* KO (*Tent5d^-/null^*); partial infertility in both male and female *Tent5a* KO (*Tent5a^-/-^*) animals; complete infertility in *Tent5b/c* double KO (dKO; *Tent5b^-/-^ Tent5c^-/-^*) females (with one wild-type allele of either gene sufficient for normal fertility). Surprisingly, the presence of GFP tag at the C-terminus of TENT5B (TENT5B-GFP) also impacts fertility - homozygous *Tent5b*^gfp/gfp^ females are completely infertile, and even *Tent5b^wt/gfp^* have reduced fertility. In contrast, no such effect is observed for N-terminally tagged TENT5B (GFP-TENT5B).

Despite partial infertility of *Tent5a* KO, male sperm and testis morphology is normal, and we observed several cases of fertile *Tent5a* KO males. In females, ovarian morphology (Supplemental Figure 1. C) and histological structure (Supplemental Figure 1. D) remain normal, and they produce a healthy pool of oocytes at the GV stage (Supplemental Figure 1. E). Furthermore, isolated GV oocytes can be matured into metaphase II oocytes *in vitro* (Figure 1. C). Taken together, these results show that the subfertility of *Tent5a* KO animals is not directly related to the process of gametogenesis itself but rather to the previously reported^29^ poor general condition of animals.

Thus, we conclude that TENT5C and TENT5D are essential for spermatogenesis, whereas TENT5B and TENT5C play a crucial but redundant role in oogenesis.

### Lack of TENT5B and TENT5C leads to early oogenesis arrest

As mentioned previously, a complete double KO of *Tent5b* and *Tent5c* results in female infertility, contrasting the lack of phenotypes in mice with the disruption of *Tent2* (*Gld2*)^18^. Notably, even one wild-type allele of either *Tent5* gene is sufficient to maintain normal fertility, pointing to their redundant roles. Accordingly, both genes are highly expressed in GV oocytes, as observed in the oocytes of *Tent5c^GFP^*and *Tent5b^GFP^* knock-in females expressing respective C-terminal GFP fusions (Figure 2. A). When we examined 12-week-old *Tent5b/*c dKO females, we detected markedly abnormal ovaries compared with heterozygous and wild-type (WT) littermates. They were smaller and showed no signs of significant follicular growth or earlier ovulation (Figure 2. B, Supplemental Figure 2. A). Neither of those could be restored by hormonal stimulation with pregnant mare’s serum gonadotropin (PMSG) and human chorionic gonadotropin (hCG).

**Figure 2.**
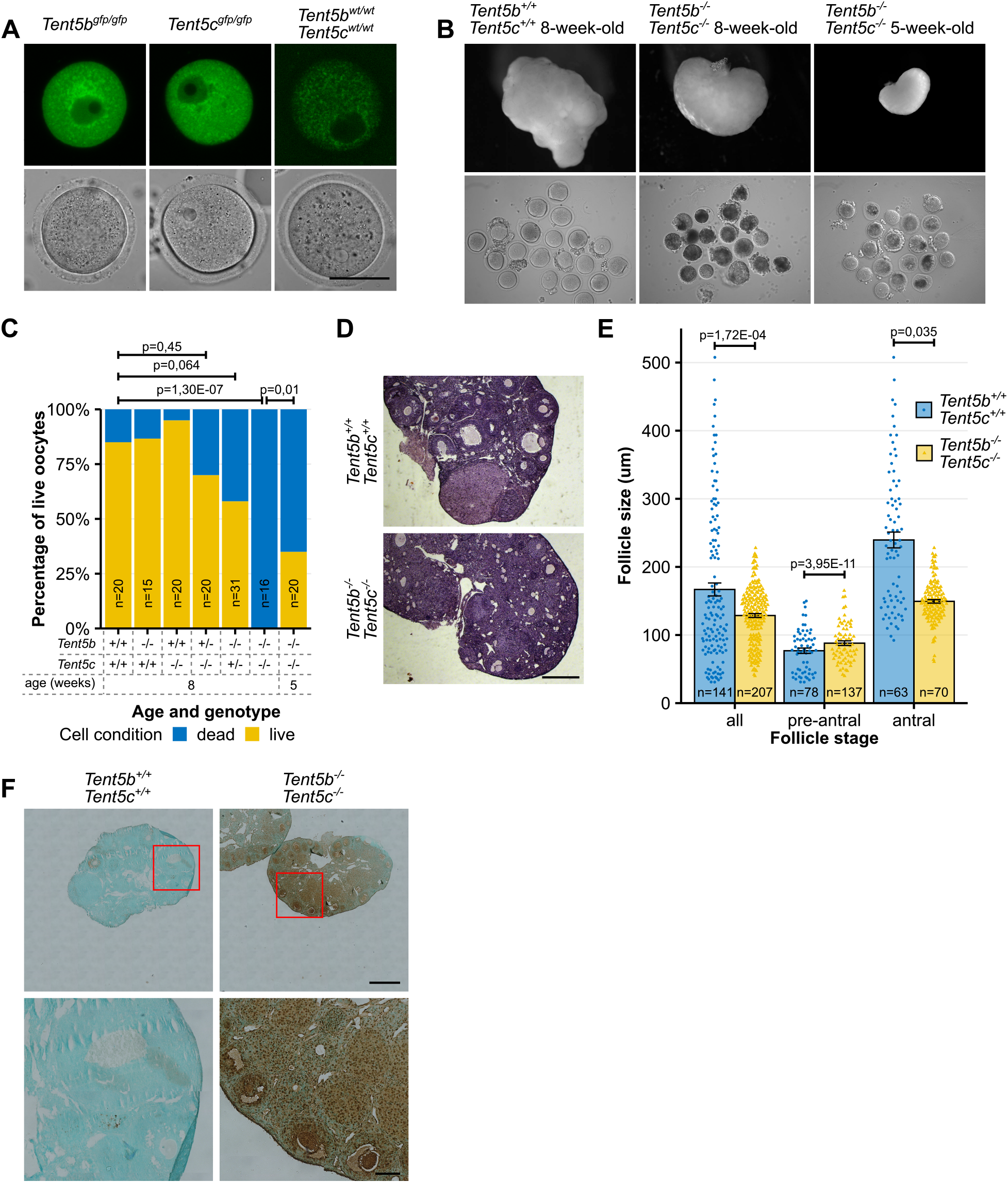
Lack of TENT5B and TENT5C leads to early oogenesis arrest. **A.** Expression pattern of TENT5B-GFP and TENT5C-GFP in GV oocytes visible as fluorescence of GFP. Scale bar = 50 µm. **B.** Morphology of ovaries and GV oocytes isolated from them in adult (8-weeks-old) and young (5-weeks-old) females. Ovaries pictures not in scale. See also Supplemental Figure 2. A. **C.** Percentage of live and dead GV oocytes isolated from the ovaries of *Tent5b/c* dKO mouse line females of different genotypes. N values represent number of oocytes counted; p-values reported for comparison of live to dead oocytes ratio in Fisher exact test, two-tailed. **D.** Haematoxylin and eosin staining of *Tent5b^+/+^ Tent5c^+/+^* and *Tent5b^-/-^ Tent5c^-/-^* ovaries’ histological sections. Scale bar = 500 µm. **E.** Comparison of ovarian follicle size distribution in *Tent5b^+/+^ Tent5c^+/+^* and *Tent5b^-/-^ Tent5c^-/-^*females, with all oocytes analysis broken down into pre-antral and antral stage follicles. Size value represents the mean width of the follicle in the widest point and width perpendicular to that measurement in narrowest point, measured on each follicle’s widest cross section available. Individual data points and n values represent individual follicles measured; bars represent mean value; error bars represent SEM; p-values reported for follicle size comparison in Mann-Whitney-Wilcoxon test. **F.** TUNEL assay staining for signs of apoptosis in ovaries’ histological sections of WT and *Tent5b/c* dKO females. Upper scale bar = 500 µm, lower scale bar = 100 µm. See also Supplemental Figure 2. B.

Dissection of *Tent5b/*c dKO ovaries revealed that all oocytes from 8-week-old females were in various stages of degeneration and cell death. In contrast, WT females of the same age maintained a significantly high (85%) population of live oocytes. In comparison, ovaries from single KO females (*Tent5b^-/-^* or *Tent5c^-/-^*) contain a pool of viable oocytes comparable to that of WT females, while females with single wild-type allele of either gene (*Tent5b^+/-^c^-/-^, Tent5b^-/-^c^+/-^*) appear to have slightly lower number of living oocytes, but they also do not differ significantly from WT females. To verify whether this was a congenital or acquired condition, we also dissected 5-week-old female ovaries. In these, oocytes showed the first signs of degeneration as seen in adult females, with 35% of oocytes remaining in the living state and the remainder showing various abnormalities - a result that differed significantly from both WT and 8-week-old *Tent5b/c* dKO females (Figure 2. B, C).

Histological staining of individual sections of the ovaries confirmed the arrest of follicular growth in *Tent5b/c* dKO females at the early antral stage. We found neither mature antral follicles ready for ovulation nor the presence of corpora lutea, indicating previous ovulation (Figure 2. D). In *Tent5b/c* dKO females, the observed follicle diameter ranged from 39.8 µm to 227.8 µm, whereas in WT it ranged from 30.85 µm to 507.7 µm (Figure 2. E). Additionally, high levels of apoptosis detected with TUNEL assay were exclusive for *Tent5b/c* dKO ovaries. We observed apoptosis not only within oocytes but also in surrounding granulosa cells. Ovaries of wild type, *Tent5b* KO and *Tent5c* KO females showed only minor traces of positive TUNEL assay staining (Figure 2. F, Supplemental Figure 2. B), completing the picture of entire *Tent5b/c* dKO ovaries deterioration – morphological, physiological, and cellular.

### Dominant effect of TENT5B C-terminal GFP knock-in

A model that provided us with additional insight into the role of TENT5B and TENT5C in oocyte biology was mouse knock-in line expressing C-terminally GFP tagged TENT5B (TENT5B-GFP), which, as we already stated, displays specific infertility of females. At the same time, N-terminal tagging of TENT5B (GFP-TENT5B) or C-tagging of TENT5C (TENT5C-GFP) did not lead to any abnormalities in the oocytes. This corresponds with GFP expression patterns in oocyte’s cytoplasm, TENT5B-GFP mean expression (measured as a level of GFP fluorescence intensity) is over three times higher than GFP-TENT5B (Figure 3. A, B).

**Figure 3.**
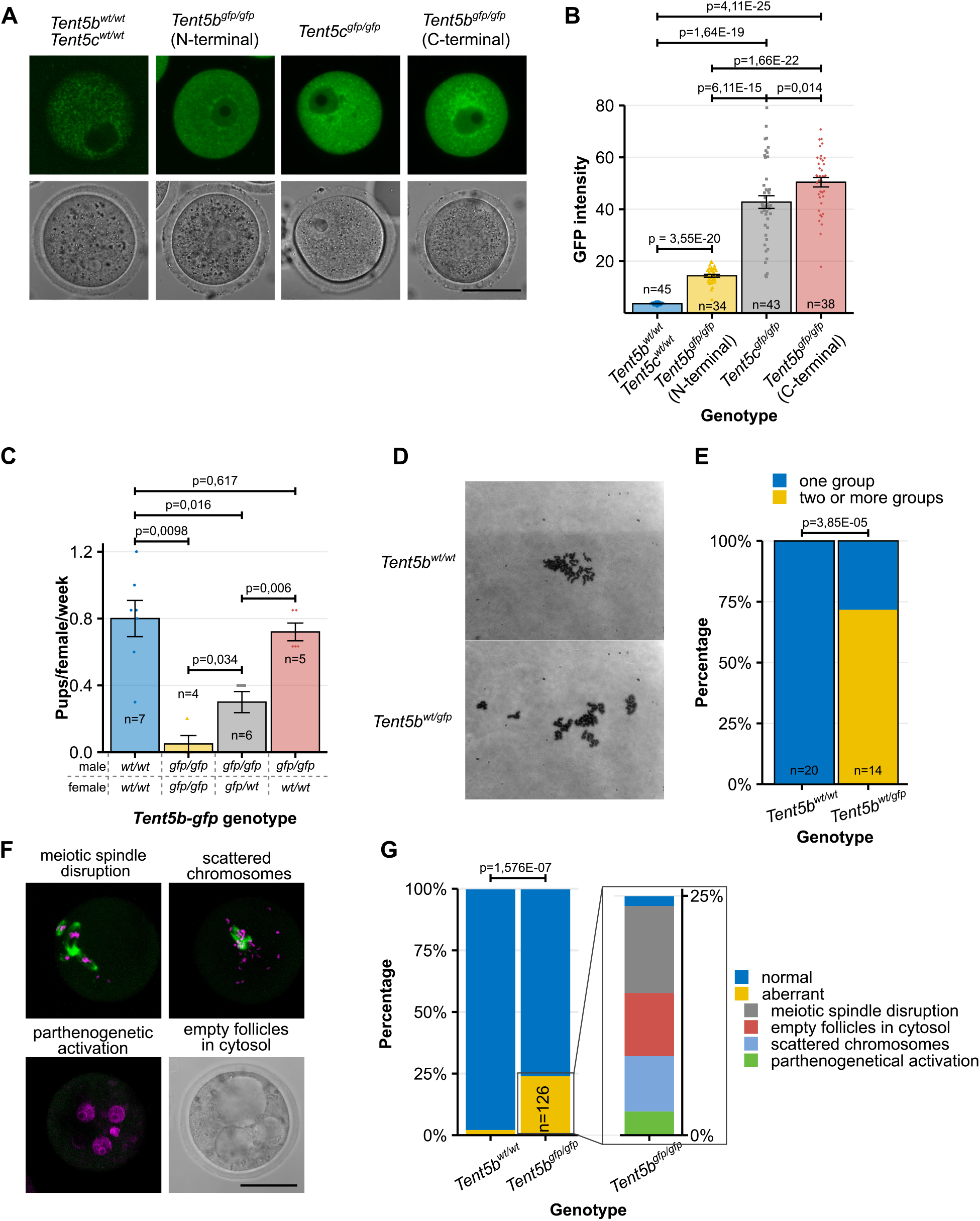
Dominant effect of TENT5B C-terminal GFP knock-in. **A.** Expression pattern of TENT5B-GFP, TENT5C-GFP and GFP-TENT5B in GV oocytes visible as fluorescence of GFP. Scale bar = 50 µm. **B.** Comparison of TENT5B and TENT5C expression levels depending on the presence of GFP tag on proteins N- and C-terminus, measured as a GFP fluorescence intensity. Individual data points and n values represent individual oocytes analyzed, bars represent mean intensity values, error bars represent SEM; p-values reported for mean values comparison in t-test, two-tailed. **C.** Birth rate for different mating configurations depending on male and female C-terminal *Tent5b^GFP^* genotype, presented as a number of pups born by individual females per week. Individual data points and n values represent individual females observed for 10 weeks, bars represent mean value, error bars represent SEM; p-values reported for number of pups comparison in Mann-Whitney-Wilcoxon test. **D-E.** Chromosome scatter preparations showing chromosome condition and organization in *Tent5b^wt/wt^* and *Tent5b^wt/gfp^* MII oocytes. N values represent number of individual oocytes analyzed; p-value reported for comparison of the ratio of oocytes with one group of chromosomes to oocytes with chromosomes separated into two or more groups in Fischer exact test, two-tailed. **F.** Chromatin (magenta) and alpha-tubulin (green) immunofluorescence staining in*Tent5b^wt/gfp^* MII oocytes. Visible are examples of different chromosomal segregation errors after oocytes ovulation. Scale bar = 50 µm. **G.** Frequency distribution of different chromosomal segregation and chromatin organization abnormalities in *Tent5b^wt/gfp^*MII oocytes. N values represent individual oocytes analyzed; p-value reported for comparison of normal to abnormal oocytes ratio in Fisher’s exact test, two-tailed.

The detrimental effect of the GFP expression at the C-terminus of TENT5B manifests as an extreme drop in litter sizes, with the severity of this effect depending on the number of GFP alleles in females, which points to a potential dominant effect. To accurately assess this effect, we set up a series of mattings in different female and male genotype configurations. We monitored them for 10 weeks, accounting for all pups. Comparing the average number of pups born by female per week (during 10-week observation), we observed that the negative effect of C-terminal GFP on fertility is limited to females. Both *Tent5b^wt/wt^* and *Tent5b^gfp/gfp^* males have a similar number of pups born when mated with WT females, which fertility gradually and significantly drops with the rising number of mutation-carrying alleles (Figure 3. C).

Simultaneously, we searched for the cause of this lowered fertility, and knowing the phenotype of *Tent5b/c* dKO females, we focused our investigation on oocyte development. Unlike *Tent5b/c* dKO, oocytes from *Tent5b^GFP^* (both *Tent5b^wt/gfp^* and *Tent5b^gfp/gfp^*) females remain alive in the ovaries and can progress further in meiosis through GV to second metaphase (MII) stage. However, chromosome spreads from those MII oocytes isolated from heterozygous (at the time homozygous *Tent5b^GFP^* females were born at very low rate and were unavailable for these experiments) and control WT females revealed the first signs of the issue underlying observed infertility (Figure 3. D). Although all oocytes of both groups maintained a standard number of 20 pairs of unaltered chromosomes, in the majority of knock-in oocytes analyzed, either one or several chromosomes were separated from the main group of chromosomes, while all chromosomes were gathered together in all WT oocytes (Figure 3. E). The reason for this was revealed by immunofluorescence staining of chromatin and spindle microtubules in MII *Tent5b^wt/gfp^* oocytes, of which one-fourth suffered some sort of disruption, most notably spindle and chromosomes disorganization, but also parthenogenetic activation and empty follicles filling cytosol (compared to only 2,38% in WT oocytes; Figure 3. F, G).

Together with the fact that *Tent5b* KO mutation alone does not impact fertility in any way, our results strongly suggest the dominant-negative or gain of function effect of GFP tag located at the C-terminus of TENT5B and the importance of TENT5B activity in the oocyte.

### TENT5s polyadenylates mRNAs in oocytes, which tight regulation is essential for oogenesis

As described above, TENT5B and TENT5C play essential roles in oogenesis. To identify the effect of dysfunction of these poly(A) polymerases at the mRNA level and identify their potential substrates, we performed Direct RNA Sequencing (DRS) using nanopore-based MinION platform (Figure 4. A, Supplemental Table 2). DRS enables genome-wide poly(A) tail length profiling and does not suffer from PCR amplification-dependent biases, which can be particularly pronounced for long poly(A) tails often present in developing gametes. The limitation of this methodology is that it requires a relatively large quantity of material for sequencing, rendering sequencing of RNA from groups of oocytes impossible in our setup. Therefore, we instead have used total RNA isolated from whole ovaries of WT, *Tent5b/c* dKO, as well as homozygous and heterozygous *Tent5b^GFP^* 30-day-old females.

**Figure 4.**
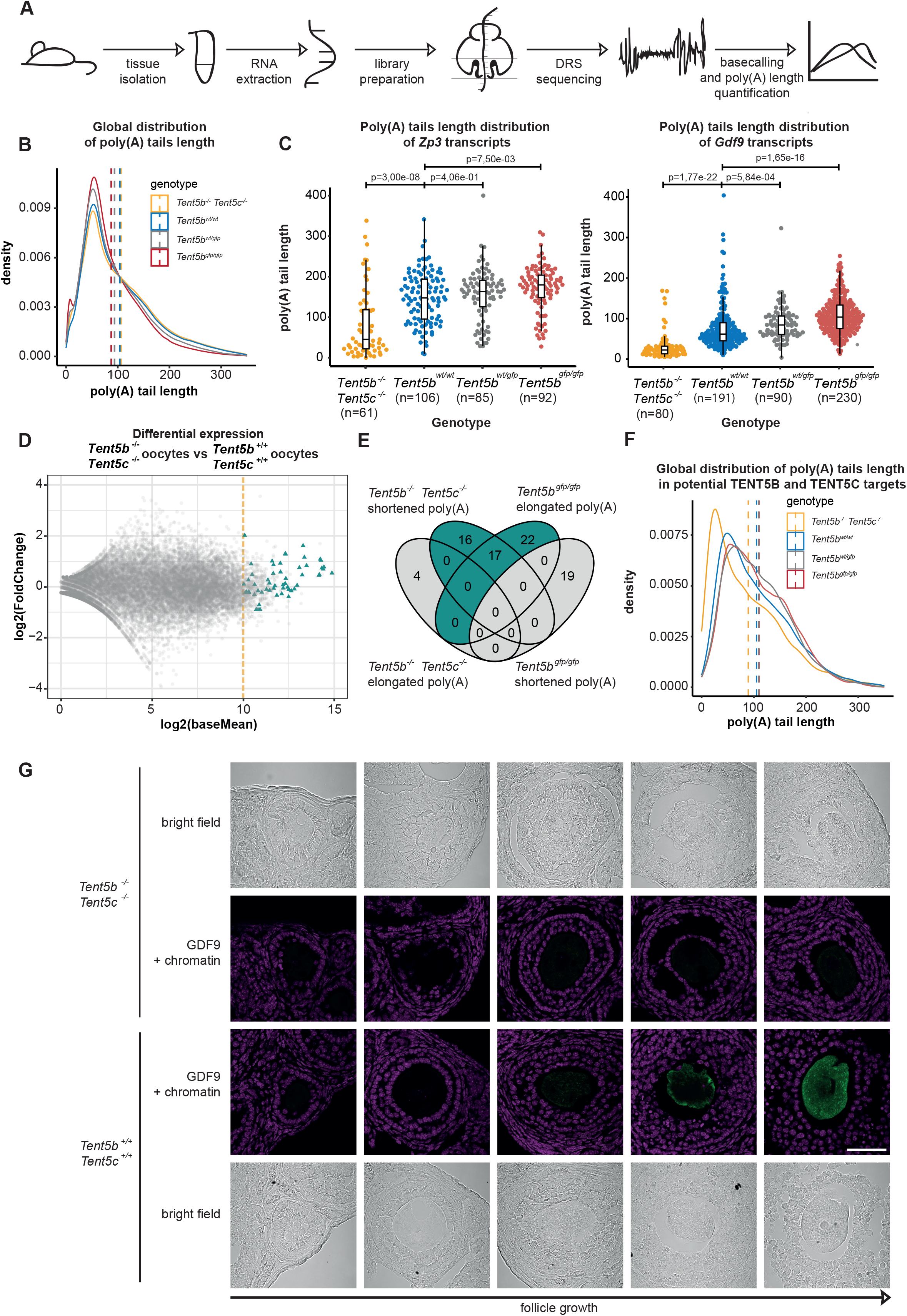
TENT5s polyadenylates mRNAs in oocytes, which tight regulation is essential for oogenesis. **A.** Schematic of the DRS-based poly(A) length profiling. **B.** The global distribution of poly(A) tail lengths of RNA isolated from *Tent5b^-/-^ Tent5c^-/-^*, *Tent5b^wt/wt^*, *Tent5b^wt/gfp^* and *Tent5b^gfp/gfp^* mice. Dashed lines indicate the mean poly(A) tails lengths: *Tent5b^-/-^ Tent5c^-/-^* – 106, *Tent5b^wt/wt^* - 103, *Tent5b ^wt/gfp^*- 93, *Tent5b ^gfp/gfp^* - 87. **C.** DRS-based poly(A) tail lengths profiling of *Zp3 and Gdf9* mRNAs isolated from ovaries. Median poly(A) tail lengths for *Zp3*: *Tent5b^-/-^ Tent5c^-/-^* =49 nt; *Tent5b^wt/wt^* = 141 nt; *Tent5b ^wt/gfp^* = 156 nt; *Tent5b^gfp/gfp^* = 170 nt; p-values reported for comparison in Mann-Whitney-Wilcoxon test with Bonferroni Hallberg correction. Median poly(A) tail lengths for *Gdf9*: *Tent5b^-/-^ Tent5c^-/-^* =28 nt; *Tent5b^wt/wt^* = 66 nt; *Tent5b ^wt/gfp^* = 78 nt; *Tent5b^gfp/gfp^* = 106 nt; p-values reported for comparison in Mann-Whitney-Wilcoxon test with Bonferroni Hallberg correction; individual data points represent individual reads analyzed **D.** RNAseq performed on oocytes isolated from *Tent5b^-/-^ Tent5c^-/-^*and *Tent5b^wt/wt^* mice. Yellow dashed line illustrates accepted cut-off that revealed 522 genes. Most potential TENT5B/C targets, highlighted by triangles are highly expressed in *Tent5b^-/-^ Tent5c^-/-^* oocytes. Differential expression calculated using DESeq2. **E.** Venn diagram illustrating the overlaps in sets of transcripts with statistically significant changes in the poly(A) tail length when compared with Mann-Whitney-Wilcoxon test, alpha=0.05 **F.** Changes in the global distribution of poly(A) tail lengths for group of transcripts selected as potential TENT5B and TENT5C targets. Dashed lines indicate the mean poly(A) tails lengths. **G.** Immunohistochemistry staining of GDF9 (green) and chromatin (magenta) in ovaries of *Tent5b^-/-^ Tent5c^-/-^* and *Tent5b^+/+^ Tent5c^+/+^* females. Scale bar = 50 µm.

*Tent5b/c* dKO had essentially no effect on global mRNA poly(A) tail distribution in ovaries (mean lengths 104 in WT vs 106 in *Tent5b/c* dKO; Figure 4. B). However, differential poly(A) tail lengths distribution analysis revealed in *Tent5b/c* dKO samples several mRNAs with significantly shorter tails encoding proteins involved in oogenesis, like mRNA encoding Zona Pellucida Glycoprotein 3 (ZP3) or Growth Differentiation Factor 3 (G*DF*9) (Figure 4. C, Supplemental Figure 3, Supplemental Table 3). At the same time, many of these had elongated poly(A) tails in homozygous and heterozygous *Tent5b^GFP^*knock-in mice (Figure 4. C, Supplemental Figure *3*,Supplemental Table 4), consistent with the fact that C-terminal GFP tag leads to TENT5B protein accumulation. Curiously, in homo- and heterozygous *Tent5b^GFP^* knock-in ovaries, we observed global shortening of poly(A) tails (mean lengths 94 and 87, respectively) which most probably represents a secondary effect (Figure 4. B). To look more specifically at oocyte-enriched transcripts, we performed an RNA-seq analysis on oocytes isolated from 5-week-old WT and *Tent5b/c* dKO females on the Illumina platform (Figure 4. D). Analysis of poly(A) tails of 522 genes highly expressed in oocytes (Supplemental Table 5) revealed 33 mRNAs that had shortened poly(A) tails in *Tent5b/c* dKO and 39 with elongated tails in *Tent5^gfp/gfp^*(Figure 4. E). Notably, 17 mRNAs were common between these groups (Figure 4. E). Combined poly(A) profile of all 72 mRNAs responding to TENT5 dysfunction (Figure 4. F) revealed an opposite effect of KO and TNETB-GFP knock-in supporting the hypothesis that GFP at the C-terminus of TENT5B leads to gain-of-function phenotype. It further indicates that mRNAs with elongated tails in *Tent5^gfp/gfp^* or shortened in *Tent5b/c* dKO represent direct TENT5s targets (Figure 4. D).

Interestingly, differential expression analysis performed on WT and *Tent5b/c* dKO Illumina dataset indicates no downregulation of genes subjected to poly(A) tail shortening in *Tent5b/c* dKO. Nonetheless, immunohistochemistry on GDF9 revealed the drastic reduction of the protein level in *Tent5b/c* dKO ovaries (Figure 4. G), pointing to the previously suggested dominant role of poly(A) tail dynamics in the regulation of translation during gametogenesis.

In sum, TENT5B and TENT5C polyadenylate a group of mRNAs essential for oogenesis, which enhances their expression. Notably, both lack of TENT5 mediated poly(A) tail extension and its excess lead to aberrant oogenesis.

### TENT5C and TENT5D are essential for different stages of spermatogenesis

While neither *Tent5* gene is single-handedly responsible for undisturbed oocyte development, this is not the case for spermatogenesis, since, as already mentioned, both *Tent5c* and *Tent5d* KO mutations render males sterile. However, the expression of both proteins is markedly different in the testis and developing sperm. Immunofluorescence detection of TENT5C-GFP and TENT5D-GFP in testis’ histological sections shows high expression of *TEN*T5C close to the lumen of seminiferous tubules in both round and elongated spermatids for both 3- and 8-week-old males. At the same time, TENT5D displays a rather uniform expression across testis’ tissue in young males, with limited amounts in earlier meiotic stages, and fades from the lumen in adult males (Figure 5. A).

**Figure 5.**
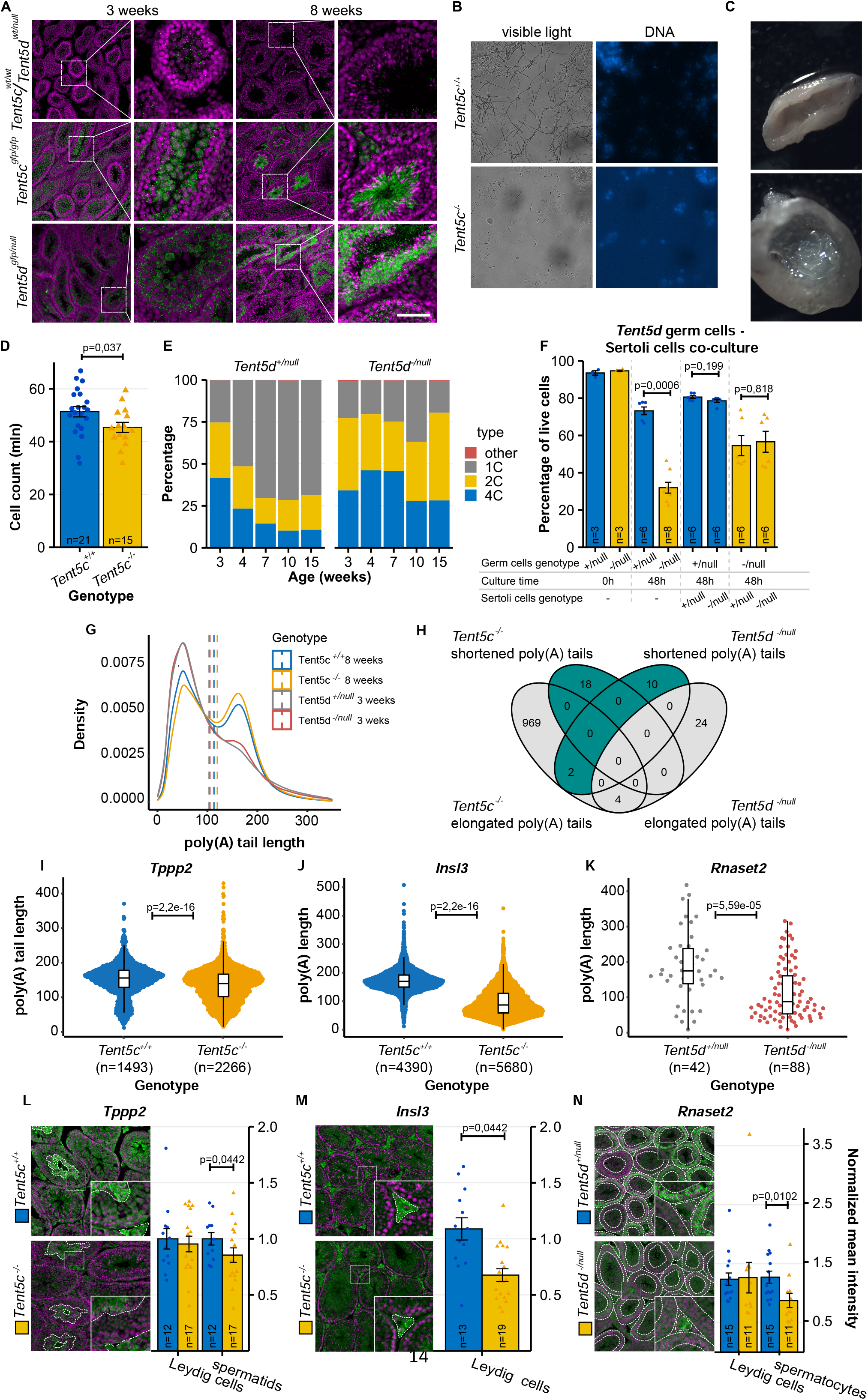
TENT5C and TENT5D are essential for different stages of spermatogenesis. **A.** TENT5C-GFP and TENT5D-GFP expression in 3- and 8-week-old testes visualized through immunohistochemistry staining of GFP (green) and chromatin (magenta). Scale bar = 100 µm. **B.** DNA staining of *Tent5c^+/+^* and *Tent5c*^-/-^ epididymis-isolated sperm. **C.** Morphology of 28-week-old *Tent5d^- /null^* males’ testes which became ‘hollow’ inside due to severe tissue degeneration. **D.** Total germ cell count (in millions) isolated from *Tent5c^+/+^*and *Tent5c^-/-^* males’ testes. Individual data points and n values represent individual males from which germ cells were isolated, bars represent mean value and error bars represent SEM; p-value reported for mean comparison in t-test, two-tailed. **E.** Changes in distribution of germ cells with different DNA content (representing different spermatogenesis stages) in *Tent5d^+/null^*and *Tent5d^-/null^* testes during the first 15 weeks of males life (single male was used for each timepoint). **F.** Germ cell survival in 48h of *in vitro* culture depending on *Tent5d* genotype of germ cells and presence and genotype of Sertoli cells. Individual data points and n values represent cell culture of germ cells isolated from single male, bars represent mean values, error bars represent SEM; p-values reported for comparison in Mann-Whitney-Wilcoxon test. **G.** The global distribution of poly(A) tails lengths of RNA isolated from *Tent5c^+/+^*, *Tent5c^-/-^*, *Tent5d^+/null^*, *Tent5d^-/null^* mice. Dashed lines indicate the mean poly(A) tails lengths: *Tent5c^+/+^* – 113, *Tent5c^-/-^* - 119, *Tent5d^+/null^* - 103, *Tent5d^-/null^* - 104. **H.** Venn diagram illustrating the overlaps in sets of transcripts with statistically significant changes in lengths of the poly(A) tails, Mann-Whitney-Wilcoxon test, alpha =0.05. **I-J.** Changes in lengths of the poly(A) tails in transcripts essential for male gametes development. DRS-based poly(A) lengths profiling of *Tppp2* and *Insl3* mRNAs isolated from *Tent5c^+/+^* and *Tent5c^-/-^*testis. Median poly(A) lengths for *Tppp2*: *Tent5c^+/+^* =152 nt; *Tent5c^-/-^* = 136 nt; individual data points represent reads analyzed; Mann- Whitney-Wilcoxon rank sum test were calculated; Median poly(A) tail lengths for *Insl3*: *Tent5c^+/+^* =169 nt; *Tent5c^-/-^* = 86 nt; Mann-Whitney-Wilcoxon rank sum test were calculated. **K.** Changes in poly(A) tail lengths in transcripts essential for male gametes development. DRS-based poly(A) lengths profiling of *Rnaset2* mRNAs isolated from *Tent5d^+/+^* and *Tent5d^-/-^*testis. Median poly(A) tail lengths for *Rnaset2*: *Tent5d^+/+^*= 175 nt; *Tent5d^-/-^* = 88 nt; individual data points represent reads analyzed; Wilcoxon rank sum test were calculated. **L-N.** Immunohistochemistry staining of chromatin (magenta) and proteins (green) encoded by transcripts identified as possible TENT5C (panels L, M) and TENT5D (panel N) substrates in testes cross-sections. Individual data points and n values represent mean values of all ROIs from individual specimen, bars represent mean values, error bars represent SEM, p-value reported for Mann-Whitney-Wilcoxon test.

To check the exact pattern of TENT5C and TENT5D expression, we sorted germ cells isolated from TENT5C and TENT5D GFP and FLAG tag knock-in males into different spermatogenesis stages based on DNA content and chromatin structure (Supplemental Figure 4) and used them for western blot (FLAG tag lines) and flow cytometry (GFP lines). Obtained results confirmed that TENT5C is initially expressed in secondary spermatocytes (spcII) with high protein levels maintained throughout the rest of spermiogenesis process, while TENT5D expression, present already in primary spermatocytes, peaks in spcII cells to rapidly drop in a round and elongated spermatids (RS and ES) (Supplemental Figure 5. A-D)

Expression patterns of TENT5C and TENT5D correspond with sperm phenotypes that we and others ^32,33^ observed – *Tent5c* KO mutation causes a high rate of head or DNA material loss in epididymis isolated sperm (Figure 5. B), while in *Tent5d* KO males’ testes become progressively smaller through mice life compared to WT males (Figure 5. C, Supplemental Figure 5. E), a result of severe tissue structure deterioration, rendering testes of 28-week-old males “hollow” inside (Figure 5. C). In consequence, the blood testosterone level of *Tent5d* KO males was significantly lower than in WT males (Supplemental Figure 5. F).

To obtain deeper insight into the spermatogenesis process we performed a more detailed flow cytometry analysis of cell cycle. *Tent5c* KO males produce on average fewer total germ cells than WT males (Figure 5. D) but accumulate more elongated spermatids (ES) in seminiferous tubules’ lumen (Supplemental Figure 5. G). *Tent5d* KO mutation hinders germ cells progress through the meiosis – while in WT males 4C population of primary spermatocytes diminishes in mice life in favor of a rapidly growing number of 1C cells (spermatids and spermatozoa), in KO mice this process is absent, and germ cell populations ratio remains stable through 15 weeks of mice life (Figure 5. E).

Then, we aimed to determine whether changes in germ line alone cause abnormal germ cell development and survivability or also depend on surrounding somatic cells. To this end, we co-cultured WT, *Tent5c* KO and *Tent5d* KO germ and Sertoli cells in different configurations for 48h and accounted for live cells after that time. In case of both genes, KO germ cells exhibit higher mortality than WT cells regardless of the Sertoli cells presence in culture. Culturing germ cells, regardless of their genotype, with Sertoli cells improved their survivability. However, this effect was independent of the genotype of Sertoli cells. Mortality and its improvement in the environment of Sertolli cells were particularly visible in the case of *Tent5d* KO cells (Figure 5. F) and only to a lesser extent for *Tent5c* KO (Supplemental Figure 5. H).

All results described above and the differential TENT5s localization indicate that the phenotypes are mainly related to germ cells and that the TENT5C and TENT5D substrates in the testis may differ. To identify these substrates, we have performed DRS of mRNA isolated from the testes of adult *Tent5c* KO and 3-week-old *Tent5d* KO males (due to their age-dependent testis degeneration). *Tent5c* and *Tent5d* KO mutations both had only minor effects on the global poly(A) tail profile (Figure 5. G, Supplemental Table 6) which may be partially attributed to changes in cell populations. At the same time, the age of mice has quite a dramatic effect as the mean poly(A) tail length in 3-week-old was 103 and increased to 113 in the adults. Importantly, we identified dozens of potential TENT5 substrates with significantly shorter poly(A) tails, with almost no overlap between mRNAs identified in *Tent5c* and *Tent5d* KO samples (Figure 5. H, Supplemental Table 7, 8). Among mRNAs affected by TENT5 dysfunction in sperm, particular attention is owed to several that play important roles in gametogenesis: *Tppp2*, *Insl2* (in *Tent5c* KO) and *Rnsset2* (in *Tent5d* KO) (Figure 5. I-K). Quantitative immunochemistry analysis revealed that shortening of their mRNA poly(A) tails leads to decreased expression at the protein level at spermatogenesis stages corresponding to KO phenotypes: TPPP2 in *Tent5c* KO spermatids and RNASET2 in *Tent5d* KO spermatocytes. Additionally, INSL3 expression was lowered in *Tent5c* KO Leydig cells (Figure 5. L-N).

### TENT5s enhance the expression of secreted proteins during gametogenesis

Poly(A) tail profiling using DRS allowed us to identify substrates of TENT5 poly(A) polymerases in the testes and ovaries, many of which play essential roles during gametogenesis. This raises the question of the mechanism leading to substrate specificity. First, we searched for sequence features of mRNAs targeted by TENT5B and TENT5C in ovaries, TENT5D in young testes, and TENT5C in adult testes. However, this analysis did not reveal any strongly enriched sequence motives, and only in the case of TENT5D target some C-reach sequences were selected (Supplemental figure 6. A-C). Then, we looked for differences in more general parameters of mRNAs regulated by TENT5s. Differences in the length of UTR segments (Figure 6. A) and GC content (Figure 6. B), although statistically significant, were negligible. Interestingly, the most distinctive feature was the coding sequence length, and TENT5 targets had relatively short open reading frames (Figure 6. A). In the case of TENT5D, the difference was the most dramatic, with a mean length of 666nt in TENT5D-regulated mRNA compared to 1493nt in the rest of mRNAs considered as a background.

**Figure 6.**
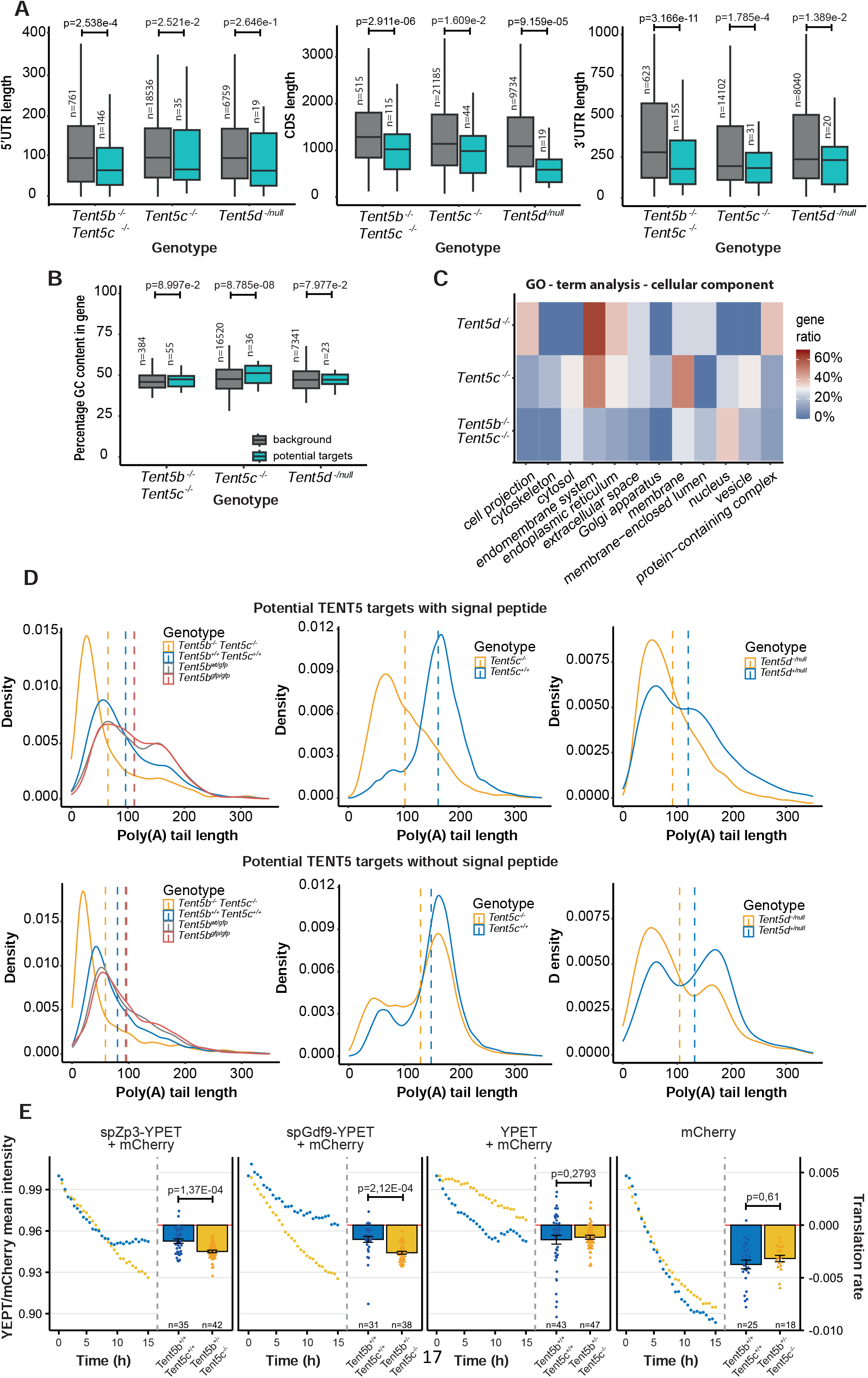
TENT5s enhance the expression of secreted proteins during gametogenesis. **A.** Differences in 5’UTR, 3’UTR and CDS length between potential targets of TENT5 proteins compared to the rest of the transcriptome detected by DRS. In the ovaries, analyses were performed for 522 oocyte-enriched mRNAs. Mann-Whitney-Wilcoxon rank sum test was calculated. **B.** Differences in GC content between potential targets of TENT5 proteins. Mann-Whitney-Wilcoxon rank sum test were calculated. **C.** Gene ontology (GO) analysis of potential TENT5 protein targets. **D.** Distribution of poly(A) tail length in potential targets of TENT5 encoding proteins with and without signal peptides. Dashed lines indicate the mean poly(A) tails lengths: *Tent5b^-/-^ Tent5c^-/-^* - 64, *Tent5b^wt/wt^* - 96, *Tent5b ^wt/gfp^* - 111, *Tent5b ^gfp/gfp^* - 111 *Tent5c^-/-^* - 102, *Tent5c ^+/+^* - 162, *Tent5d^-/-^* - 91, *Tent5d ^+/+^* -120; for substrates with signal peptide and *Tent5b^-/-^ Tent5c^-/-^* - 58, *Tent5b^wt/wt^* - 80, *Tent5b ^wt/gfp^* - 95, *Tent5b ^gfp/gfp^* - 96, *Tent5c^-/-^* - 129, *Tent5c ^+/+^* - 148, *Tent5d^-/-^* - 104, *Tent5d ^+/+^* - 132 for substrates without signal peptide. P-values for TENT5B/C substrates with and without signal peptide were calculated by Mann-Whitney-Wilcoxon test with Bonferroni Hallberg correction, and all p<2.2e-16. P-values for TENT5C and TENT5D substrates with and without signal peptide were calculated by Mann-Whitney-Wilcoxon rank sum test, and all p<2.2e-12 **E.** Normalized mean intensity of fluorescence and calculated translation rate of oligoadenilated reporter GFP mRNAs with signal peptide injected into *Tent5b^+/+^ Tent5c^+/+^* and *Tent5b^+/-^ Tent5c^-/-^*GV oocytes. Mean intensity data points represent mean YFP fluorescence intensity normalized to fluorescence of control polyadenylated reporter mRNA encoding mCherry. Individual translation rate data points and n values represent individual oocytes analyzed; bars represent mean values, error bars represent SEM; p-values reported for t-test, two-tailed.

Since the sequence analysis did not provide any clue about the potential mechanism of TENT5s substrate selection, we moved on to a more functional analysis. Unsurprisingly, Gene Ontology (GO) analysis revealed genes involved in reproduction and the developmental processes as the most enriched terms (Supplemental Figure 6. D-E). Moreover, a noticeable fraction of TENT5 substrates represent mRNAs encoding secreted proteins (Figure 6. C). In testes, such mRNA represents the majority of TENT5 substrates. The search for the endoplasmic reticulum-targeting signal peptide confirmed this view (Figure 6. D). In testis, the signal peptide was found in 8 out of 10 proteins coded by TENT5D substrates and 9 out of 18 TENT5C substrates. This fraction was smaller in ovaries where 13 of 55 TENT5B and TENT5C substrates encoded proteins with a signal peptide. Notably, in the case of both testes and ovaries effect of TENT5s dysfunction on the lengths of poly(A) tails were stronger for mRNAs encoding proteins with ER targeting signal than for the rest of the affected mRNAs (Figure 6. D).

Finally, to answer the question of whether the presence of a signal peptide is determining factor of TENT5s mediated regulation in the oocyte, we injected GV oocytes with mRNA reporters encoding signal peptide of either GDF9 or ZP3 (oocyte specific TENT5s targets essential for oogenesis) followed by YPET fluorescent protein-coding sequence and short sequence coding ER-retention KDEL motif. Each signal peptide mRNA reporter had a short oligo-A tail of 20 adenines at the 3’-end to facilitate further polyadenylation. Along with such reporters, every oocyte was also injected with *in-vitro* polyadenylated, mCherry-encoding mRNA to control translation without the need for such a particle to be polyadenylated in the cell. This way, the dynamics of YPET fluorescence intensity, normalized to mCherry intensity, directly represent the effect of polyadenylation of the reporter mRNA translation in individual oocytes. For this experiment, we chose to use *Tent5b^+/-^ Tent5c-/-* as they remain alive in ovaries but may already display changes at the transcriptomic level. Indeed, YPET reporters with a signal peptide have a lower translation rate in *Tent5b^+/-^ Tent5c^-/-^* oocytes compared to WT ones. When the YPET reporter lacks the ER-targeting signal, the translation rate remaines unchanged between WT and *Tent5b^+/-^ Tent5c^-/-^* oocytes – similarly as for the oocytes that were only injected with mCherry reporter as a control of the translation process itself (Figure 6. E).

## Discussion

In this study, we have demonstrated the essential roles of TENT5B, C and D poly(A) polymerases during gametogenesis in mice. In spermatogenesis, in addition to TENT5C and TENT5D, previous studies identified TPAP, a testis-specific poly(A) polymerase, which is present in the cytoplasm of spermatogenic cells. Its knockout also leads to male infertility^34^. Consequently, three distinct cytoplasmic poly(A) polymerases are essential for proper spermatogenesis. In contrast, in oogenesis there is a redundancy between TENT5B and TENT5C targeting overlapping pools of transcripts.

Although the mechanism of substrate recognition by TENT5s is far from being understood, it becomes apparent that mRNAs encoding proteins targeted to the endoplasmic reticulum represent the major group of TENT5 substrates in testis and ovaries. This result aligns with the function of TENT5s in the enhancement of expression of secreted immune effector proteins in B cells and macrophages^31^, constituents of extracellular matrix essential for bone mineralization by osteoblasts^29^ or secreted antimicrobial peptides in the gut of the worm *C. elegans*^31^. Among TENT5B and TENT5C substrates are secreted and oocyte-specific zona pellucida components (ZP2 and ZP3) and signaling protein GDF9, both essential for proper oocyte development and fertility. While early studies of murine *Zp3* KO and null mutations showed that its essential for zona pellucida (ZP) formation and oocyte’s ovulation along cumulus complexes^35,36^, recently described Zp3 mutations in human patients were showed to lead to Empty Follicle Syndrom (EFS), where no oocytes can be isolated from patients’ ovarian follicles due to oocytes degeneration or failure to develop properly^37,38^. GDF9 is known to be expressed exclusively in the oocyte^39^, but its role extends beyond it, affecting granulosa cells proliferation^40,41^ and regulating development of entire ovarian follicle^42,43^, with its deficiency phenotype resembling our observation in *Tent5b/c* dKO female mice. TENT5C in testis is essential for proper expression of TPPP2, a protein vital for sperm motility and its fertilization capacity, and gonad tissues-specific hormone INSL3, expressed in Leydig cells^44^, where its indispensable for testicular descent^45,46^ and promotes germ cell survival^47^. Finally, TENT5D controls the expression of extracellular ribonuclease RNASET2 involved in the innate immune response ^48^,tumor suppression^49^ and, most importantly for this work, regulation of sperm motility^50,51^. *R*na*set2* KO mice were also shown to recapitulate neurological degeneration symptoms of human RNASET2 deficiency^52^.

Interestingly, there are differences between the effect of TENT5-mediated polyadenylation in somatic cells and during gametogenesis. TENT5 proteins in B cells, macrophages, and osteoblasts mainly regulate mRNA stability. Their dysfunctions always lead to decreased levels of its substrates through the half-life regulation. The same is for completely synthetic mRNA, such as the Moderna mRNA-1273 vaccine, which is taken by macrophages after intermuscular injection. Vaccine mRNA encoding coronavirus Spike protein is targeted to ER and recruits TENT5A for polyadenylation and stabilization^53^. Consequently, TENT5A is essential for efficient antigen production and immune responses. In contrast, during gametogenesis, TENT5 KO leads to shortening its substrates’ poly(A) tails, leading to decreased levels of proteins produced, while mRNA levels, however, remain unchanged. This agrees with previously proposed poly(A) tail metabolism differences between somatic cells compared to gametes and early embryos^54^. In the latter, there are limiting amounts of the poly(A) binding proteins essential for efficient translation. As a result, poly(A) tail extension enhances protein synthesis at the same time in oocytes, and during spermatogenesis mRNA stability is regulated differently than in somatic cells.

It has been known for years that maintaining the balance of poly(A) tail in transcripts accumulated in oocyte’s cytoplasm during its development is crucial for further maturation and fertilization. Until the discovery of TENT5s, the main known mechanism postulated for this regulation of mRNA poly(A) tails in oocytes was the CPEB protein accompanied by a complex of proteins organized around it^23^. CPEB recognized specific motifs in mRNA sequence and was postulated to recruit multiple different effectors like PARN deadenylase and the GLD2 poly(A) polymerase to maintain a short oligo(A) tail of the transcript. When the hormonal stimuli arrive to the oocyte, promoting its maturation, PARN is supposed to dissociate from the complex, allowing GLD2 to elongate the transcript’s poly(A) tail, activating it. The unchanged fertility of *Gld2* KO undermines this model and may suggest that TENT5s act redundantly with GLD2^18^. However, the data provided by us indicate that TENT5s act at the stage of oocyte growth when transcription is active rather than during oocyte activation. Furthermore, in the TENT5 substrates, there are no CPEB binding sites. Thus, the poly(A) polymerase or polymerases responsible for waves of cytoplasmic polyadenylation during oocyte activation remain to be established. All these reveal previously unexpected complexity of the regulation of poly(A) tail dynamics during gametogenesis in mammals.

## Author contributions

M.B and A.D. wrote the final version of the manuscript with contributions from A.C.-C., O.G., and M.K.-K., M.B. performed TENT5B and TENT5C expression measurements in oocytes, phenotype analysis of *Tent5a* KO, *Tent5b/c* dKO and *Tent5b^GFP^* mice, GDF9 immunohistochemistry staining, oocyte mRNA reporter injection and prepared oocyte and ovary RNA sequencing libraries, M.B. and M.S. performed *Tent5a* KO oocyte maturation, O.G. performed blood analysis, body weight measurements and *Tent5c* and *Tent5d* phenotype analysis, M.K.-K. performed germ cells – Sertoli cells co-cultures, all flow cytometry experiments and cell sorting, M.S. performed *Tent5b^GFP^* mating experiment and chromatin analysis experiments, B.T. and K.J. performed all immunochistochemistry experiments in testes, S.M. prepared sperm and testes RNA sequencing libraries, J.G. and E.B. established all mouse lines, M.B. reviewed and analyzed all gathered data except sequencing results, A.C.-C. supported by P.K. analyzed all RNA sequencing results, M.B. and A.C.-C. prepared figures.

## Acknowledgments

We thank Andrzej Dziembowski lab members for their support, prof. Marco Conti (UCSF) for sharing YPET and mCherry mRNA reporter-carrying plasmids, Aleksandra Brouze for critical and proofreading of the manuscript and all members of Genome Engineering Unit of International Intitute of Molecular and Cell Biology for maintenance of animal colony and animal genotyping. NGS was performed thanks to Genomics Core Facility CeNT UW (RRID:SCR_022718), using NovaSeq 6000 platform financed by Polish Ministry of Science and Higher Education (decision no. 6817/IA/SP/2018 of 2018-04-10).

This project has received funding from the European Union’s Horizon 2020 research and innovation programme under grant agreement No 810425.

The research leading to these results was funded by the Norwegian Financial Mechanism 2014 -2021, no. UMO-2019/34/H/NZ3/00733.

This project was also funded by National Science Centre, Poland, grant no. 2019/33/B/NZ2/01773.

## STAR Methods

## RESOURCE AVILABILITY

### Lead contact

Correspondence and requests for reagents and plasmids should be directed to lead contact, Andrzej Dziembowski

### Materials availability

Plasmids, mouse lines and reagents generated in this study are available at request.

### Data and code availability

Data sets used for plot generation (except RNA sequencing data) are available as Supplemental Table 9.

Raw sequencing data (fast5 files) were deposited at ENA (MinION sequencing; project accession number: PRJEB63526) and GEO (Illumina sequencing; project accession number: GSE239661) repositories and are publicly available as of the date of publication. Project accession numbers as listed in the key resources table. Additionally, ENA sample accession numbers together with DRS run details are listed in Supplemental Table 2.

This paper does not report original code.

Any additional information required to reanalyze the data reported in this paper is available from the lead contact upon request.

## EXPERIMENTAL MODEL AND STUDY PARTICIPANT DETAILS

All experiments were performed using mouse lines generated using CRISPR/Cas9 method in C57BL/6/Tar x CBA/Tar mixed background.

In this study females of following genotype and age were used: *Tent5a^-/-^* (8-12 weeks), *Tent5b^-/-^c^-/-^* (4-12 weeks), *Tent5b^-/-^c^+/-^* (8-12 weeks), *Tent5b^gfp/gfp^*(C-terminal) (8-12 weeks), *Tent5b^gfp/gfp^* (N-terminal) (4-12 weeks), *Tent5b^gfp/wt^* (N-terminal) (4-12 weeks), *Tent5c^gfp/gfp^*(8-12 tygodni), wild-type (4-12 tygodni), and males of following genotype and age were used: *Tent5b^gfp/gfp^*(8-48 weeks), *Tent5c^-/-^* (8-48 weeks), *Tent5d^null/-^* (3-48 weeks), *Tent5c^gfp/gfp^*(8-48 weeks), *Tent5c^flag/flag^* (8-48 weeks), *Tent5d^gfp/gfp^* (3 weeks), *Tent5d^flag/flag^* (8-48 weeks), wild-type (3-48 weeks).

The mice were kept in conventional polystyrene cages in the animal facility of the Faculty of Biology, University of Warsaw. A 12/12 light cycle was maintained in the room, 15 air exchanges per hour. The relative humidity in the rooms was 55% ± 10%. The temperature in the rooms was 22°C ± 2°C. During all procedures and breeding the animals received rodent feed and ad libitum water.

Sex of animals and cells used was determined by the according fertility phenotype connected to particular mutations.

## METHODS DETAILS

### Mouse line generation

Generation of *Tent5a* KO and *Tent5c* KO mouse lines was described previously^29,30^. Procedure for generation of *Tent5a* KO, *Tent5b* KO, *Tent5d* KO, GFP-*Tent5b, Tent5b*-GFP, *Tent5c-*GFP, *Tent5c-*FLAG, *Tent5d-*GFP and *Tent5d-*FLAG mouse lines is described below. Chimeric sgRNAs were synthesized by T7 RNA Polymerase *in-vitro* transcription and purified by PAGE. Cas9 mRNA was *in-vitro* transcribed with T7 RNA Polymerase (produced inhouse) and subsequently polyadenylated using *E. coli* Poly(A) polymerase (NEB, Cat# M0276L) and m7Gppp5’N Cap was added using Vaccinia Capping System (NEB, Cat# M0280S). For knock-in mouse lines, repair DNA template was purchased as double stranded gene fragments (GeneArt, Thermo Fischer Scientific). See Supplemental Table 1. for sgRNAs, repair templates and genotyping oligonucleotides sequences.

Zygotes obtained from mated females were microinjected into the cytoplasm using Eppendorf 5242 microinjector (Eppendorf-Netheler-Hinz GmbH) and Eppendorf Femtotips II capillaries with the following CRISPR cocktail: Cas9 mRNA (25 ng/µl), sgRNA (15 ng/µl) and donor dsDNA (7,5 ng/ µl) After overnight culture microinjected embryos at 2-cell stage were transferred into the oviducts of 0,5-day p.c. pseudo-pregnant females. Born pups were genotyped by PCR at around 4 weeks. The presence of mutation was confirmed by sequencing in the founder mouse and, after backcrossing, in N1 generation mice. See Supplemental File 1. for primers’ sequences used for genotyping.

### Cell isolation and culture

#### Oocyte and embryo

In order to stimulate ovarian follicle growth and ovulation, adult female mice (aged 8-12 weeks) were injected intraperitoneally with pregnant mare’s serum gonadotropin (PMSG, Folligon, Intervet) and human chorionic gonadotropin (hCG, Chorulon, Intervet) respectively (10 units of hormone in 100 µl of PBS) in 44-46h interval.

All procedures and culture of germinal vesicle (GV) and metaphase-II (MII) stage oocytes as well as zygotes, unless stated otherwise, were performed in pre-warmed M2 or M16 medium (Sigma, Cat# M7161 and Cat# M7292), additionally supplemented with 150 ng/µ l dbcAMP (Sigma, Cat# D0260) to prevent meiosis resumption of GV prophase I-arrested oocytes when needed. For oocyte isolation, hormonally stimulated females were sacrificed either 48h after PMSG stimulation for GV oocytes or 16h after hCG stimulation for MII oocytes. For zygote isolation, hCG stimulated females were mated with males overnight and checked for presence of vaginal plug following day as the insemination indicator and then sacrificed 21-22h post hCG injection. Ovaries and oviducts were dissected and immediately placed in M2 culture medium. For GV oocytes isolation, cumulus-oocyte complexes were released to medium by follicle puncture and cumulus cells surrounding oocytes were removed by pipetting. For MII oocytes and zygote isolation, oviducts were moved to dish containing warmed Hyaluronidase solution (300 µg/ml in M2, Sigma, H4272) facilitating cumulus cells detachment, cumulus cells-surrounded oocytes/zygotes were released by ampulla puncturing and pipetted to disperse cumulus cells. Before and between further procedures, denuded oocytes were cultured in 10 µl drops of M2 medium under mineral oil in groups of 10-20 on plastic dishes (35x10 Tissue Culture Dishes, Falcon) placed in incubator at 37°C and 5% CO2 in the air.

#### Cell isolation and culture

The protocol used for male germ cells isolation from testis was adapted from the work of Bastos *et al.*^55^. Dissected testes were decapsulated and placed in Falcon with 25 ml of HSBB buffer (Gibco, Cat# 14025-092) at room temperature. Next, 1 ml of collagenase (12.5 mg/ml) was added (Sigma, Cat# C-7657) and tubes were put to shake at 32°C for 20 minutes in the shaking water bath (120 osc/min). Tubes were manually agitated every 5 minutes to facilitate the dissociation of the tubules. The tubules were washed once in 25 ml 1x HSBB buffer and resuspended in 10 ml of the HSBB buffer with 200 µl of the stock trypsin (25 mg/ml, Sigma, Cat# T-6763) and 2 µl of the stock DNAse I (5 mg/ml, Sigma, Cat# DN-25) and again put in a water bath with agitation at 32°C for 15 minutes (120 osc/min). After the incubation, samples were dissociated for 4 minutes by pipetting 10 ml serological pipet and centrifuged at 1500 rpm for 3 min. The supernatant was discarded, and cells were resuspended in 1 ml of HSBB with 10% FBS (Gibco, Cat# 10270106) and counted in Thoma chamber. Prepared cells were used for further procedures.

Isolation of Sertoli cells from testis was based on the protocol published by Bhushan *et al.* and Anway *et al*.^56,57^. Dissected testes were decapsulated and digested with 10 ml of Trypsin (2.5 mg/ml)-DNase I (10µg/ml) solution (Sigma, Cat# T5266) for 4 min in water bath 32°C (120 oscilations/min). Next the trypsin digestion was stopped by adding 5 ml trypsin inhibitor (10 mg/ml) (Sigma, Cat# T6522), samples were vigorously mixed and incubated for 5 min. After tubules settled down in the tube supernatant was carefully removed and 10 ml of trypsin inhibitor (2,5 mg/ml) was added. Digested tissue was washed 9 times to remove all the germ cells. Remaining seminiferous tubules were digested by Collagenase (1 mg/ml)-Hyaluronidase (1 mg/ml; Sigma, Cat# H3506)-DNase I (10 µg/ml) mixture followed by Hyaluronidase (1 mg/ml)-DNase I (10 µg/ml) digestion. Finally digested seminiferous tubules were passed 10 times through the 18G needle. Isolated Sertoli cells were then seeded in the concentration of 1 x 10^6^ cells/ml in RPMI medium (Gibco, Cat# A10491-0) supplemented in 10% FBS and cultured in 37°C, 5% CO2 until a monolayer of cells was obtained.

Previously isolated germ cells were seeded in the concentration of 2 x 10^6^ cells/ml onto the Sertoli monolayer and cultured for 48h. After 48h medium containing germ cells were taken, and germ cells were washed with HSBB buffer and prepared for the flow cytometry analysis.

### Morphology, histology, cell visualization

#### Tissue and cell fixation

For tissue histology analysis, H&E staining and TUNEL assay tissues were fixed in 10% Neutral Buffered Formalin (Sigma, Cat# HT501128) for 24h, dehydrated in ethanol dilution series (70% x1, 95% x2, 100% x3, changes every hour with overnight incubation in 4^th^ change of 100% ethanol; Chempur, Poland) and 2 changes of xylene, and incubated in two changes of liquid paraffin followed by paraffin embedding on metal trays using EC 350 Tissue Embedding Center (Myr). Paraffin-embedded tissues were sectioned using semi-automatic microtome (Leica RM2125 RTS) to obtain 10 µm-thick tissue slices transferred on Superfrost Ultra Plus glass slides (Thermo Fisher Scientific, Cat# 11976299) and dried in 37°C overnight before further processing.

For immunohistochemistry analysis tissues were fixed in 4% paraformaldehyde (Sigma, Cat# P6148) in 0.1 M phosphate buffer for 24h and cryopreserved in two changes of 30% sucrose (Merck, Cat# S0389) in 0.1 M PB, 24h each. Tissues were frozen in Killik medium (Bio-Optica, Cat# 05-9801), cut into 20 µm-thick sections using cryostat and placed on Superfrost Ultra Plus glass slides. Sections were then stored in 4°C until further processing.

Oocytes for immunofluorescence were fixed in 4% PFA (Thermo-Fisher Scientific, Cat# 28908), permeabilized with 0.5% Triton-X100 (Sigma, Cat# T9284), 30 min in RT each, and blocked with 3% BSA (Sigma-Aldritch, Cat# A7906) in 4°C overnight.

#### H&E staining

For nucleus-cytoplasm staining, paraffin-embedded sections on glass slides were first deparaffinized in two changes of xylene (10 minutes each) and rehydrated in 5 changes of ethanol (100% x2, 96% x2, 70% x1, 2 minutes each) and distilled water for 2 minutes. Subsequently, slides were stained in Harris Hematoxilin (Kolchem, Poland) for 10 minutes, rinsed for two minutes in tap water, dipped once in acidic ethanol (1% HCL solution in 70% ethanol), again briefly rinsed in tap water and for 2 minutes in Tap Water Substitute (2.5g NaHCO_3_ and 20g MgSo_4_ * 7H_2_O in 1 liter of water) and finally stained in Eosin Y (Kolchem, Poland) for 1 minute. After that, slides were dehydrated by reversing the rehydration protocol above (with 5 minutes xylene incubation instead of 10), and sealed using DPX mounting medium (Sigma, Cat# 44581) and a cover slide, and dried overnight.

#### TUNEL assay

Tissue sections on glass slides where deparaffinized, rehydrated and sealed as described above for H&E stained and a commercially available TUNEL Assay Kit (Abcam, Cat# ab206386) was used according to the manufacturer’s protocol to detect double stranded breaks of DNA created during apoptosis. Negative control staining was prepared by substituting TdT in reaction mixture with water and positive staining control was prepared by treating sections with 1µg/µl DNase I in TBS (tris-buffered saline) for 20 min.

#### Oocyte immunofluorescence

MII oocytes were incubated overnight at 4°C in primary monoclonal antibody against β-tubulin conjugated with FITC (Sigma-Aldritch, Cat# F2043; 1:50) and then washed twice in PBS for 15 min. To visualize chromatin, oocytes were stained in droplets of propidium iodide (0.01 mg/ml in PBS) at 37.5 °C, for 30 min on glass bottom dishes (MatTek Corp.).

#### Chromosome spreads

MII oocytes were treated by Acidic Tyrode’s Solution (Sigma, Cat# T1788) for about 30 s at 37°C to remove zona pellucida. After wash in M2 medium oocytes were placed for 1-5 minutes in hypotonic solution of 1% sodium citrate. Next, single oocyte was placed on standard glass slide and fixed by several drops of methanol/acetic acid (3:1). Slides were air dried and chromosomes were stained with 4% Giemsa solution for 10 minutes. Slides were washed in distillated water, air dried, washed in xylene and finally mounted using DPX mounting medium.

#### Immunohistochemistry

Frozen sections were dried for 10’ at 55°C, washed twice for 10’ in TBS, incubated for epitope retrieval in pressure-cooker (70-90kPa, 160°C) for 2’ in 20mM Tris Buffer, pH 9,0 (for Insl3 and Tppp2), pH 7,5 (for Rnaset2) or 10mM citrate buffer, pH 6,0 (for Gdf9), and washed twice briefly with TBS. Next, sections were incubated 1h at room temperature (RT) in blocking buffer (10% serum from the species that subsequently used secondary antibody was raised in, 1% BSA and 0,3% Triton X-100 diluted in TBS) and then overnight at 4°C in moisture chamber with primary antibodies (anti-Tppp2 1:100, AbCam, Cat# ab236887; anti-INSL3 1:200, Invitrogen, Cat# PA5-55921; anti-RNASET2 1:100, Sigma-Aldrich, Cat# HPA029013; anti-GDF9, Abcam, Cat# ab254323) diluted in the same buffer. Next day slides were washed 3x for 5’ with TBS-T (TBS with 0,025% Triton X-100), incubated 1h at RT with secondary antibodies (donkey anti-rabbit Alexa 568-conjugated, 1:500, Invitrogen, Cat# A-10042 or for GDF9 staining goat anti-rabbit Alexa586-conjugated, 1:500, Invitrogen, Cat# A-11011) diluted in blocking buffer with Hoechst 33342 (Sigma, Cat# H3570) 10µg/ml, washed 3x for 5’ with TBS-T and once with TBS.

To quench autofluorescence all sections except GDF9 staining were incubated with TrueBlack Plus(R) (Biotium, 23014) diluted 1x in PBS for 10’, washed 3x for 5’ with PBS. All sections were sealed with cover slide using ProLong Gold Antifade Mountant (Invitrogen, Cat# P36934).

### Flow Cytometry analysis

In the seminiferous tubules of adult mammalians, germ cells in different maturation steps coexist, with 1C (round spermatids, elongating and elongated spermatids, spermatozoa), 2C (several types of G1 spermatogonia, secondary spermatocytes) and 4C (different stages of primary spermatocytes, G2 spermatogonia) DNA content^58^. For the efficient isolation of germ cells, we used protocol established by Bastos *et al.* ^55^. Considering the various sizes, shapes, and DNA content we employed fluorescence-activated cell sorting (FACS) as a convenient method to analyze cells at different stages of spermatogenesis, resulting in differentiating 5 populations for downstream analyses: gonial cells (gonia), several stages of primary spermatocytes (4C), secondary spermatocytes (spcII), round spermatids (RS), elongated spermatids (ES).

For cell cycle analysis germ cells were stained with Vybrant™ DyeCycle™ Violet Stain (Thermo Fisher Scientific, Cat# V35003) 1 µl per 1 mln of germ cells^59^. Cells were incubated at 32°C in a water bath with agitation for 35 minutes (osc 90/min). Fluorescence was excited by 405 nm laser and DCV Blue fluorescence was detected with 450/50 filters while DCV Red fluorescence was detected with 525/50 filters. For the live/dead staining, the LIVE/DEAD™ Fixable Near-IR Dead Cell Stain Kit (Thermo Fisher Scientific, Cat# L34976) was used. Fluorescence of GFP was excited by 488 nm laser and detected with 530/30 filters. Samples were analyzed with BD LSRFortessa™ and sorted with BD Aria Fusion™ under FACS Diva Software v8.0.1 (BD) software control and analyzed using FlowJo (Data Analysis Software v10)^55,59^.

### Imaging

Immunohistochemistry results were scanned as tile arrays on LSM800 (Zeiss) confocal microscope with 20x air objective. Fluorescence intensities were extracted from images using Fiji (Is Just ImageJ) ^60^ with the help of Grid/Collection Stitching Plugin (Preibisch *et al.*, 2009), manually indicated ROIs and self-written macros.

Oocyte live imaging was performed on Opera Phenix High-Content Screening System at 38°C and 5% CO_2_ in the air. Oocytes were placed in M16 medium on 384-well plate, one cell per well and were scanned for 15h in 30 minutes intervals in brightfield, YFP and mCherry channels with 5 images gathered in z-axis with 40x water immersion objective every 10 µm with germinal vesicle centre as a middle slice. Fluorescence intensity values were gathered from maximum intensity projection of all z-stack images for each oocyte using dedicated Opera Phenix software.

### Western blotting

For western blot analysis, whole tissue or equal amount of cells were lysed with 0.1% NP40 in PBS supplemented with mix of protease inhibitors and viscolase (A&A Biotechnology, Cat# 1010-100) for 30 min in 37 °C with 600 rpm shaking, then Laemmli buffer was added and samples were denatured for 10 min in 100 °C. Samples were separated on 12–15% SDS–PAGE gels, proteins were transferred to Protran nitrocellulose membranes (GE Healthcare), and then membranes were stained with 0.3% w/v Ponceau S in 3% v/v acetic acid to control amount of the protein on the membrane for every sample. Membranes were then incubated with 5% milk or 5% BSA in TBST buffer according to the technical recommendations of the antibodies’ suppliers for 1h followed by incubation with specific primary antibodies diluted 1:1:000 (Flag Tag, Life Technologies, Cat# PA1-984B) or 1:10,000 (GRP 94, Santa CruzBiotechnology, Cat# sc-11402) overnight in 4 °C. Membranes were then washed three times in TBST buffer, incubated with HRP-conjugated secondary antibody: anti-mouse (Millipore, Cat# 401215) diluted 1:5000 for 1 h at RT. Membranes were washed three times in TBST buffer and proteins were visualized by enhanced chemiluminescence acquired on X-ray film (Fujifilm, Cat# 47410 19289) using Clarity Western ECL Substrate (Biorad, Cat# 1705061).

### Oocyte microinjections

#### Reporter design

Original plasmids carrying mCherry- and YPET reporter-coding sequences, described before^61^, were received on courtesy of prof. Marco Conti (UCSF). mCherry sequence-carrying plasmid was used unchanged to produce mCherry reporter mRNA. KDEL (along additional sequences; not used in this work) motif with STOP codon was cloned in ORF downstream of YPET-coding sequence with SLIC method using DNA fragments produced by PCR from mouse oocyte’s cDNA (with forward primer containing KDEL motif-coding sequence)^62,63^ and signal peptide-coding sequence of Zp3 or Gdf9 were inserted between YPET-coding sequence and T7 reverse transcriptase promoter in ORF by HindIII and BshTI digestion and ligation of oligonucleotides with sticky ends homologuous to both enzymes’ cutting sites. All plasmids’ sequences were verified after cloning by Sanger sequencing. All oligonucleotide sequences are available in Supplemental Table 1.

#### Reporter mRNA synthesis

To synthesize YPET-coding mRNA reporters, plasmids were linearized by PCR using Phusion Hot Start II DNA polymerase (Thermo Fisher Scientific, Cat# F549) with universal forward primer at YFP sequence and universal reverse primer in Plat 3’UTR sequence with 20 thymine residues overhang (see Supplemental Table 1. for primer sequences). For 50 µl reaction volume 10ng of output DNA and 0,5µM final primer concentration was used. The PCR reaction was run in thermal cycler with following program: 98°C – 3 min., 30 cycles (98°C – 10s., 57°C – 15s., 72°C – 30s.), 72°C – 5 min., 4°C – hold.

PCR product was run on agarose gel to verify obtained band size and purified using spin-column based Clean-Up kit (A&A Biotochnology).

Plasmid containing mCherry reporter was linearized by restriction digestion with MunI (MfeI) enzyme. For 40 µl of reaction 2.5 µl of DNA sample was used. Digestion reaction was performed in 37°C for 2h and inactivated in 65°C for 20 min. Digestion product was run on agarose gel to verify obtained band size and purified with 1 volume of AMPure XP (Beckman Coulter, Cat# A63882) beads and eluted with nuclease-free water.

Resulting DNA fragments where transcribed *in-vitro* using homemade batch of T7 RNA polymerase with following 40 µl reaction setup for 2h in 37°C: 12.5 ng/µl DNA template, 4 µl T-buffer, 10mM MgCl_2_, 2.5mM of ATP, CTP, GTP and UTP, 1.5 µl Ribolock and 4 µl of T7 RNA Polymerase, H_2_O. mCherry reporter was additionally polyadenylated *in vitro* by mixing whole 40 µl of *in-vitro* transcription reaction with 10 µl of poly(A) buffer (0.5 M Tris-HCl, 2.5 M NaCl), 10 µl 50 nM MnCl_2_, 4 µl 100 mM ATP, 5 µl 5U/µl *E.coli* poly(A) polymerase and 331 µl H_2_O and incubating reaction in 37°C for 1h.

All mRNA reporters were then cleaned by LiCl precipitation. Briefly, reactions were brought up to 400 µl volume, 200 µl of 7 M LiCl, 50 mM EDTA mix were added and samples were incubated overnight at -20°C. Then samples were centrifuged for 20 min at 16000 g, 4°C, supernatant discarded, and precipitated mRNA washed with 1ml of 70% ethanol. After centrifuging for 3 min at 14000 g ethanol was discarded and air-dried pellet resuspended in RNase-free water.

Finally, all mRNA samples were capped using Vaccinia Capping System. 5 µg of mRNA in 30 µl was incubated for 5 min in 65 °C and 5 min in 4 °C before mixing with 4 µl of 10x capping buffer, 2 µl 10mM GTP, 2 µl 2mM SAM and 2 µl of Vaccinia Capping Enzyme and incubating for 30 min at 37 °C.

Capped mRNA was cleaned up by LiCl method as described above and resuspended in aprox. 20-50 µl H_2_O.

#### Oocyte injection

After isolation, oocytes were allowed to recover for approx. 1.5h. For microinjection, oocytes were transferred into new drops of pre-warmed medium. Microinjections were performed on Axio Observer 5 microscope (Zeiss) equipped with InjectMan 4, TransferMan 4r, CellTram 4r Oil and FemtoJet 4i (Eppendorf). Mixture of 12.5 ng/µl mCherry mRNA, 12.5 ng/µl YPET reporter and 0.05% NP-40 (NP-40 in PBS in nuclease-free water) in nuclease-free water was injected into cytoplasm with Femtotip II in minimal possible liquid volume (FemtoJet 4i set up for continuous leak, compensation pressure p_c_ = 50-150 hPa). After microinjection, oocytes were allowed to recover in culture for approx. 3h before imaging.

### RNA isolation

Total RNA from oocytes was isolated using PicoPure RNA isolation kit (Thermo Fischer Scientific, Cat# KIT0204) according to manufacturer’s protocol with following adjustments: groups of isolated oocytes were placed in 10 µl of provided Extraction Buffer in PCR tubes and incubated for 30 min in 42 °C on thermoblock. Samples were then stored in –80 °C. Before proceeding to next step, samples were pooled for total of 30 oocytes per sample and amount of provided ethanol used was adjusted depending on final sample volume. RNA was eluted with 11 µl of Elution Buffer provided. DNA was removed from final sample by DNase treatment using TURBO DNA-free Kit (Thermo Fisher Scientific, Cat# AM1907) according to manufacturer’s protocol. DNA-free RNA was stored in –80 °C.

RNA from whole tissue samples and cultured cells was isolated using TRI Reagent (Sigma, Cat# T9424) using manufacturer’s protocol. Tissues were homogenized in TRI Reagent using glass Dounce homogenizer and cell monolayers were rinsed with TRI Reagent and lysed cells were gathered for further isolation according to manufacturer’s protocol. For all RNA samples DNA was removed as described for oocyte samples above.

### cDNA synthesis

cDNA was produced using SuperScript III reverse transcriptase (Thermo Fisher Scientific, Cat# 18080093) according to manufacturers protocol with oligo(dT) priming and addition of ERCC RNA Spike-InMix (final dilution: 1:100000; Life Technologies, Cat# 4456740) for first-strand synthesis from oocyte-isolated RNA and RiboLock RNase inhibitor (40 units; Thermo Fischer Scientific, Cat# EO0382) for second strand synthesis in all samples.

Output material was purified in 1 volume of AMPure XP beads, eluted in 20 µl nuclease-free water and second strand synthesis reaction was set up by addition of following components: Second Strand Buffer (NEB, Cat# B6117S), RNase H (10 units; NEB, Cat# M0297S), *E. coli* DNA ligase (10 units; NEB, Cat# M0205S), *E. coli* DNA polymerase (50 units), dNTPs (final conc. 0.4 mM) and nuclease-free water for a final reaction volume of 50 µl. Reaction was incubated in 16°C overnight. Final cDNA output was purified in 1 volume AMPure XP beads and eluted with 10 µl of nuclease-free water.

### RNA sequencing

#### Illumina libraries - oocytes

Tagmentation reaction was performed following published protocols^64,65^ with various steps and amount of enzyme used optimized for use of homemade batch of Tn5 transposase produced in our laboratory. Briefly: 100 µM Tn5ME-A and Tn5ME-B oligonucleotides in annealing buffer (50 mM NaCl, 40 mM Tris-HCl pH = 8) were mixed 1:1 with 100 mM Tn5MErev oligonucleotide and incubated in thermocycler in 95°C for 5 min and 65°C for 5 min, and finaly cooled to 4°C and stored at -20 °C (with slow cooling between each step). Tn5 (0.25 mg/ml) was loaded with linker oliguncleotides Tn5ME-A/Tn5MErev and Tn5ME-B/Tn5MErev (see Supplemental File 1 for full sequences) by mixing of 10 µl Tn5 with 0.5 µl of both linkers (0.35 µM) and incubating for 45 min in 23°C with shaking at 350 rpm. Right before the reaction setup loaded Tn5 was diluted 10 times with nuclease-free water. 10 µl of freshly prepared tagmentation buffer (20mM Tris-HCl ph 7.5; 20 mM MgCl_2_; 50 % dimethylformamid) were mixed with 5 µl of diluted Tn5 and 5 µl od cDNA, incubated for 3 min at 55°C in preheated thermocycler and cooled to 10°C for 1 min. Reaction was inactivated by addition of 5 µl 0,2% SDS and incubation for 5 min at RT. Tagmented cDNA was purified in 1.25 volume of AMPure XP beads and eluted in 10 µl of nuclease-free water. For library amplification KAPA HiFi HotStart ReadyMix 2x (Roche, Cat# KK2602) with addition of 5% DMSO was used. 5 µl of tagmented cDNA was used for 15 µl reaction volume. Reaction was run in thermocycler with following program: 72°C – 15 min., 95°C – 30 s., 15 cycles: (98°C – 20s., 58°C – 15s., 72° – 30s.), 72°C – 3 min., 4°C - hold. Number of cycles yielding best results was determined experimentally and was further individually adjusted for each batch of prepared cDNA. Libraries were sequenced using Illumina NovaSeq 6000.

#### Nanopore libraries – ovaries and testes

Libraries for direct RNA sequencing (DRS) were prepared using Direct RNA sequencing kit (ONT, Cat# SQK-RNA002) following manufacturers protocol, with adjustment to magnetic beads clean-up steps: each time KAPA Pure Beads (Roche, Cat# KK8000) were used and RNA-bead mixture was incubated stationary on bench at RT. To improve sequencing performance and efficiency 90-150 ng of *Saccharomyces cerevisiae* or *Saccharomyces pombe* oligo(dT)-enriched mRNA was added to all samples. Sequencing experiments were performed on MinION device and Flow Cell Type R9.4.1 (ONT, Cat# FLO-MIN106D) and basecalling with Guppy 6.0.0 (ONT).

### Data analysis

#### DRS Nanopore sequencing analysis

To figure out poly(A) tail length DRS reads were mapped to the GencodeM26 reference transcript sequences^66^ using Minimap 2.17^67^ with options -k 14 -ax map-ont –secondary = no and processed with samtools 1.9 (samtools view -b -F 2320)^68^ to removed supplementary alignments and reads mapping to reverse strand. All unmapped reads were discarded from analysis. For each read length of poly(A) tail were estimated using the Nanopolish 0.13.2^69^ polya function and only reads tagged as PASS were considered in later analyses. Samples from the same condition were analyzed together since they were strongly corelated. Analyses of changes in median poly(A) tail length and mRNA abundance were performed using R. Supplemental Table 3and 6 contain the number of counts, mean, median and geometric mean poly(A) tail lengths. Genes represented by more than 20 reads, and with median poly(A) tail length difference WT:mutant less than –10 nt were selected as potential TENT5BC, TENT5C and TENT5D targets. We took genes represented by at least 20 reads and with medians calculated for poly(A) tail lengths greater by 10 nt than in the control as potential TENT5B targets in the GFP knock-ins.

#### Differential expression analysis

Reads obtained from Illumina and DRS sequencing were mapped to the mouse GRCm39 genome^70^ using Minimap 2.17, with options -k 14 -ax splice -uf, Features were assigned using Gencode M26. DRS datasets were processed using featureCounts^71^ from the subread package in the long read, strand-specific mode (-L -s 1), including only features covered by at least 20% (–fracOverlapFeature 0.2). Illumina datasets were mapped to the mouse GRCm39 genome using STAR^72^ on default program settings and processed by featureCounts in the short read mode (-p –O) including only features covered by at least 20% (–fracOverlapFeature 0.2). R software DESeq2^73^ were used to determine differences in mRNA expression levels. The shrinkage approach of DESeq2 was used to implement a regularized logarithm transformation for better visualization and ranking of genes. Supplemental Table 5 contains DESeq statistics for transcripts – Illumina reads dataset. Motif enrichment analysis

Motif enrichment analysis was performed on three sets of potential TENT5B/C/D targets (Supplemental Tables 4,7, 8.) in comparison to their backgrounds. The sets used as background contain the sequences of transcripts identified in the DRS data for WT, with more than 20 mapped reads per transcript. Fasta sequences of 3′ UTRs, CDS and 5′ UTRs of potential TENT5 substrates and their background were obtained using bedtools getfasta tool (v. 2.29.2)^74^, using bed files with coordinates downloaded from UCSC Table Browser tool (GENCODE M26 track and known gene table)^75^ and GRCm39 genome sequence. Sequence motifs enriched in 3′ UTRs, CDS and 5′ UTRs of TENT5B/C/D potential substrates were identified using the STREME tool^76^, run with options --dna --order 4.

#### Analysis of TENT5 B/C/D potential targets transcript’s structure

Comparison of the length of structural elements of transcripts were performed BioMart R library^77^. Estimation of the statistical significance of differences between the 3’UTR, CDS and 5’UTR lengths of potential TENT5 targets and background was performed using the Mann-Whitney-Wilcoxon test. As a background were used all transcripts without potential TENT5 targets, represented by more than 20 reads.. GC content analyses were performed using BioMart R library. Statistics for comparison GC content in sequences of potential TENT5s targets and backgrounds were calculated using Mann-Whitney-Wilcoxon test.

#### Signal peptides in potential TENT5 targets

The protein sequences of potential TENT5 targets were downloaded from the UniProt Release 2023_02 database^78^. They were searched for the presence of signal peptides using TargetP Gene ontology analysis^79^.

Gene ontology analysis was performed using the WEB-based GEne SeT AnaLysis Toolkit with ORA and geneontology options^80^.

### Data visualization

All data visualization was performed in R using ‘ggplot’ library^81^.

## QUANTIFICATION AND STATISTICSL ANALYSIS

### Translation rate calculation

For every oocyte imagined and analyzed, YPET and mCherry fluorescence values were collected as described above. Oocytes with missing data were discarded altogether from analysis. YPET value was divided by mCherry value for each individual oocyte and timepoint for normalization purposes. For each oocyte, linear regression model was calculated using ‘lm()’ function with ‘YPET/mCherry intensity ∼ timepoint’ model in Rstudio software, and beta regression coefficient (describing slopes direction and degree) was used as a translation rate.

### Statistical analysis

All statistical analyses were performed using Rstudio. All results (except sequencing data) were compared using two-sided t-test for normally distributed data, Mann-Whitney-Wilcoxon test for non-normally distributed data, Fischer exact test for 2x2 contingency tables and Chi-square test for bigger tables. Normallity was checked for all data sets by Pearson normality test. P-values lower or near assumed significancy value of 0,05 were reported on figures. Full p-values, number of samples/specimens analyzed, error bars are reported on figures and fully described in corresponding figure legends.

### Supplemental Information

Supplemental Figures 1 to 6 together with Supplemental Table 1 containing oligonucleotides, dsDNA, and sgRNA sequences, are available in Supplemental File.

ENA accession numbers of deposited raw MinION sequencing files are available in Supplemental Table 2.

Results of sequencing data analyses are available in Supplemental Tables 3-8 with following titles:

Supplemental Table 3. Poly(A) tail length statisitics for RNA samples from *Tent5b/c* dKO, *Tent5b^wt/gfp^* and *Tent5b^gfp/gfp^* ovaries.

Supplemental Table 4. TENT5B/C substrates identified in mice ovaries. Supplemental Table 5. Differential expression analysis in *Ten5b^-/-^ Tent5c^-/-^* oocytes.

Supplemental table 6. Poly(A) tail length statisitics for RNA samples from *Tent5c* KO and *Tent5d* KO whole testes.

Supplemental Table 7. TENT5D substrates identified in mice testes.

Supplemental Table 8. TENT5C substrates identified in mice testes.

Data used for plot generation (except RNA-seq data) are available in Supplemental Table 9.

