## Supplemental File for "Polyadenylation of mRNAs encoding secreted proteins by TENT5 family of enzymes is essential for gametogenesis in mice"

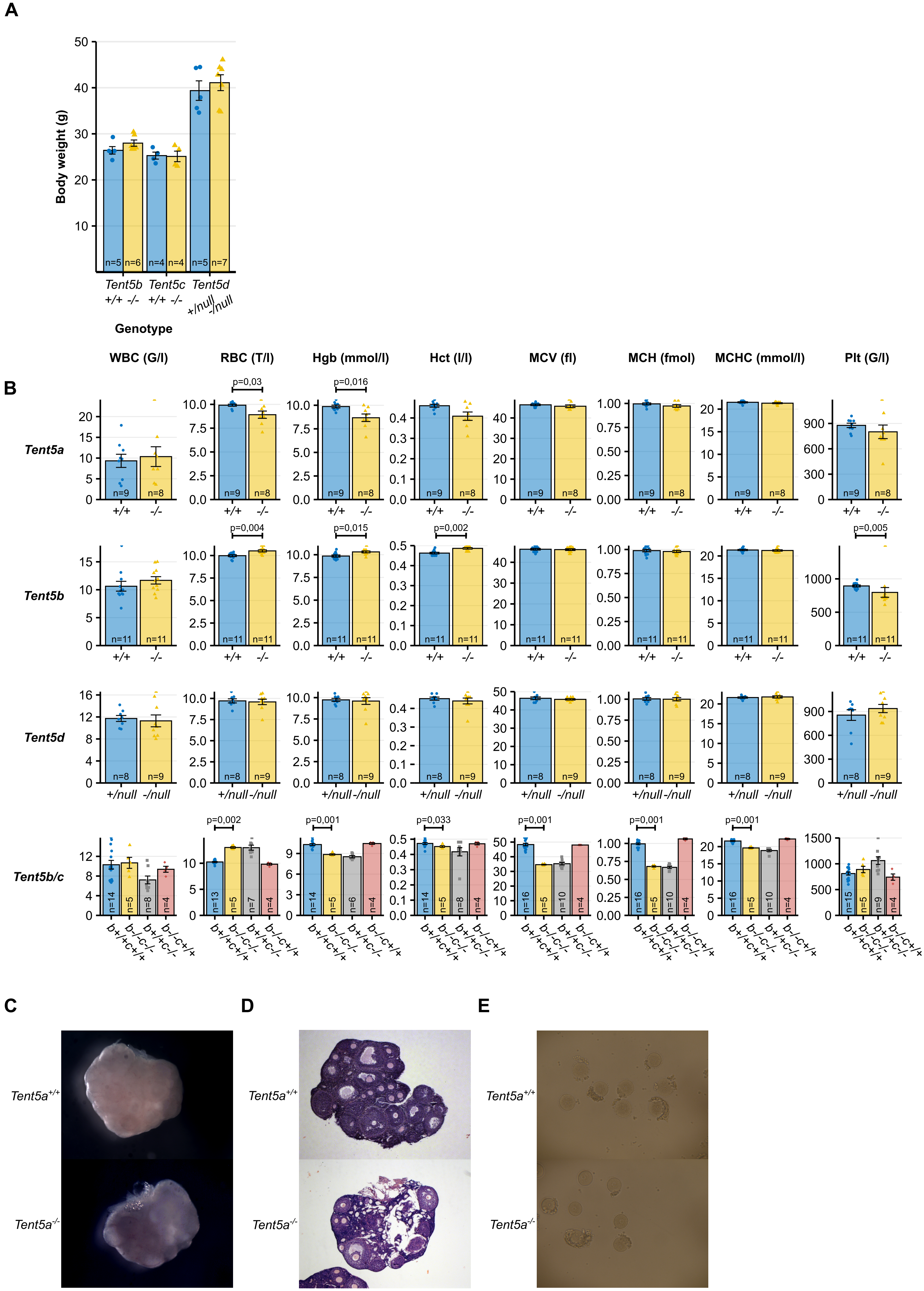
A
Body weight (g)
50
40
30
20
10
n=5 n=6 n=4 n=4 n=5 n=7
Tent5b
+/+ -/-
Tent5c
+/+ -/-
Tent5d
+/null -/null
Genotype
B
WBC (G/l)
RBC (T/l)
Hgb (mmol/l)
Hct (l/l)
MCV (fl)
MCH (fmol)
MCHC (mmol/l)
Plt (G/l)
Tent5a
p=0,03
p=0,016
n=9 n=8
+/+ -/-
Tent5b
p=0,004
p=0,015
p=0,002
p=0,005
n=11 n=11
+/+ -/-
Tent5d
n=8 n=9
+/null -/null
Tent5b/c
p=0,002
p=0,001
p=0,033
p=0,001
p=0,001
p=0,001
b+/+c+/+
b-/-c-/-
b+/+c-/-
b-/-c+/+
C
D
E
Tent5a+/+
Tent5a-/-

**Supplemental Figure 1. Basic phenotype analysis of all *Tent5* KO mice lines.**

**A.** Body weight of males of different *Tent5* KO mice lines. Animals of each mouse line were weighted at different age. Individual data points and n values represent individual animals weighted, bars represent mean values, error bars represent SEM, p-values reported for comparison in the Mann-Whitney-Wilcoxon test. **B.** Blood morphology parameters related to the *Tent5* mice lines genotype. Multiple differences in *Tent5b*$^{-/-}$ *Tent5c*$^{-/-}$ females are related to a detrimental effect of *Tent5c*$^{-/-}$ mutation alone, reported previously by Mroczek *et al.*[27]. Individual data points and n values represent blood samples from individual animals, bars represent mean values, error bars represent SEM; p-values reported for mean parameter value comparison in Mann-Whitney-Wilcoxon test; WBC = white blood cells, RBC = red blood cells, Hgb = hemoglobin, Hct = hematocrit, MCV = mean corpuscular volume, MCH = mean cell hemoglobin, MCHC = mean corpuscular hemoglobin concentration, Plt = Platelet. **C-E.** *Tent5a*$^{-/-}$ females' ovary morphology (**C**), histology (**D**) and GV oocytes (**E**) compared to *Tent5a*$^{+/+}$ ones. No changes observed in follicle growth, ovulation, and oocyte condition at GV stage.

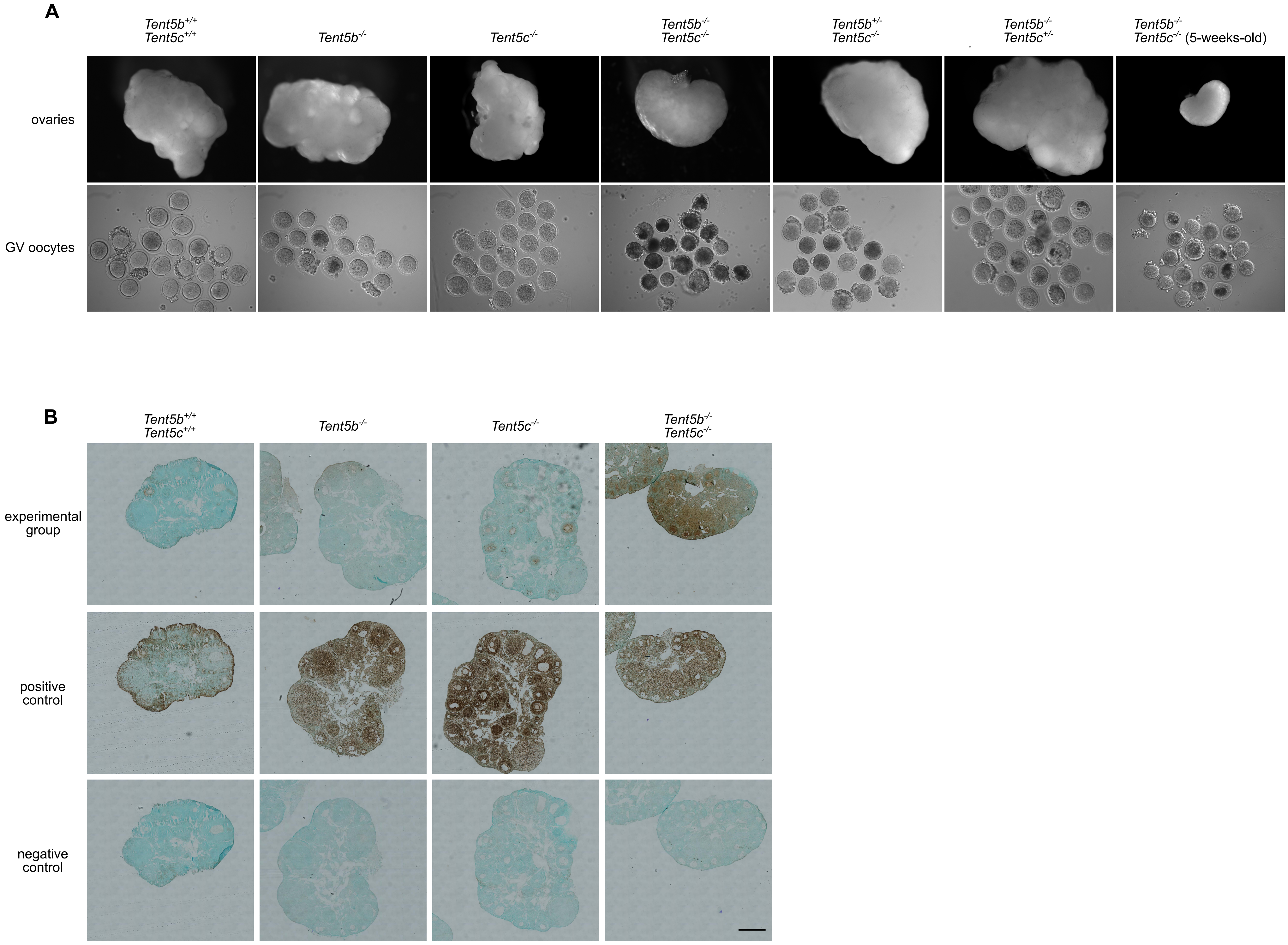
A
Tent5b+/+ Tent5c+/+
Tent5b-/-
Tent5c-/-
Tent5b-/- Tent5c-/-
Tent5b+/- Tent5c-/-
Tent5b-/- Tent5c+/-
Tent5b-/- Tent5c-/- (5-weeks-old)
ovaries
GV oocytes
B
Tent5b+/+ Tent5c+/+
Tent5b-/-
Tent5c-/-
Tent5b-/- Tent5c-/-
experimental group
positive control
negative control

**Supplemental Figure 2. *Tent5b/c* dKO ovaries and oocytes morphology and apoptosis analysis**

**A**. Morphology of the ovaries and GV oocytes of females of different *Tent5b* and *Tent5c* genotypes. Only *Tent5b*$^{-/-}$ *Tent5c*$^{-/-}$ ovaries display no signs of ovulation or follicle growth and wide oocyte degeneration. **B**. TUNEL assay for signs of apoptosis in cross-section of ovaries of different *Tent5b* and *Tent5c* genotypes. Only *Tent5b*$^{-/-}$ *Tent5c*$^{-/-}$ mice display high levels of apoptosis in both oocytes and somatic granulosa cells surrounding them.

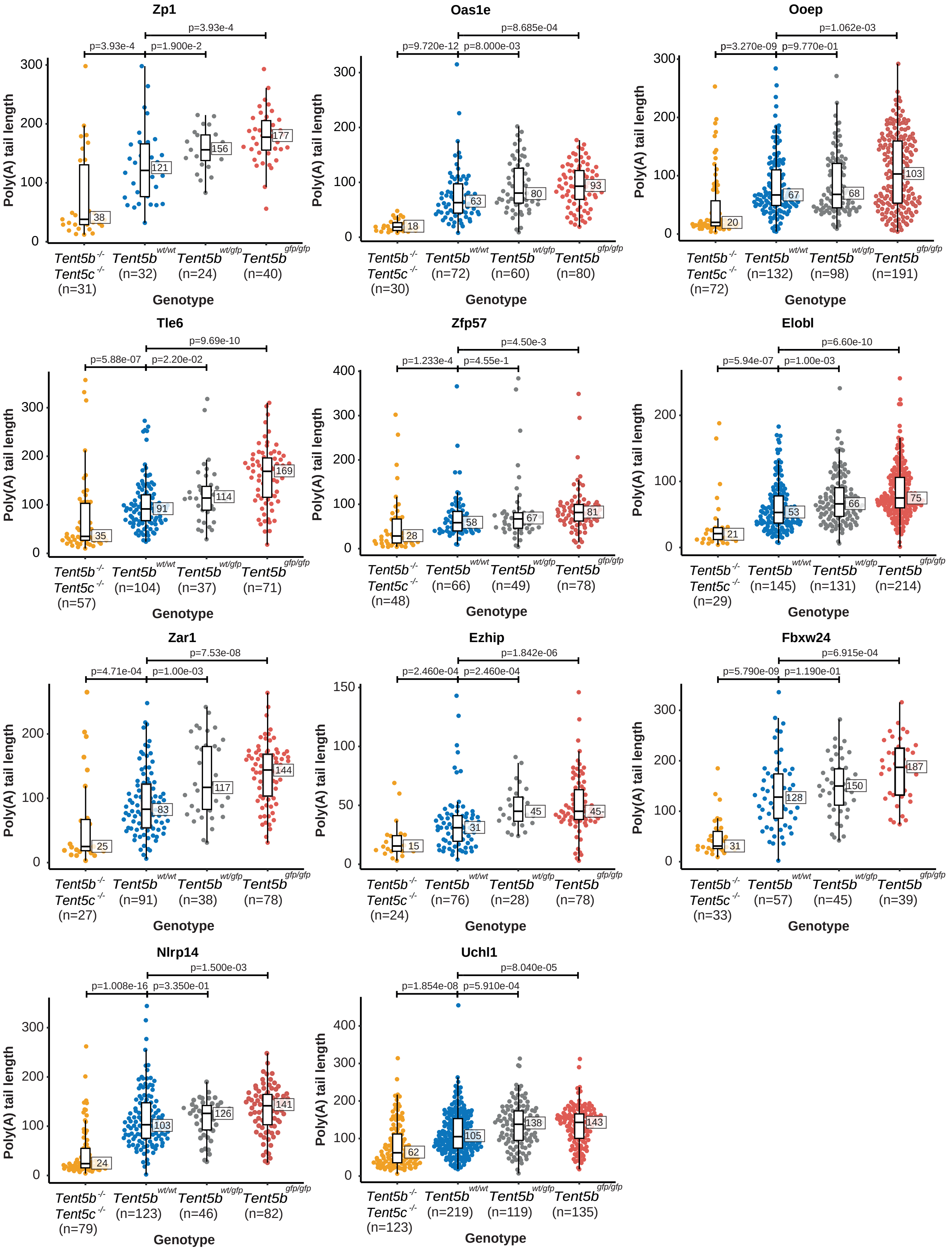

Zp1
p=3.93e-4
p=1.900e-2
p=3.93e-4
Poly(A) tail length
38
121
156
177
Tent5b-/- Tent5c-/- (n=31)
Tent5b wt/wt (n=32)
Tent5b wt/gfp (n=24)
Tent5b gfp/gfp (n=40)
Genotype
Oas1e
p=9.720e-12
p=8.000e-03
p=8.685e-04
18
63
80
93
Tent5b-/- Tent5c-/- (n=30)
Tent5b wt/wt (n=72)
Tent5b wt/gfp (n=60)
Tent5b gfp/gfp (n=80)
Ooep
p=3.270e-09
p=9.770e-01
p=1.062e-03
20
67
68
103
Tent5b-/- Tent5c-/- (n=72)
Tent5b wt/wt (n=132)
Tent5b wt/gfp (n=98)
Tent5b gfp/gfp (n=191)
Tle6
p=5.88e-07
p=2.20e-02
p=9.69e-10
35
91
114
169
Tent5b-/- Tent5c-/- (n=57)
Tent5b wt/wt (n=104)
Tent5b wt/gfp (n=37)
Tent5b gfp/gfp (n=71)
Zfp57
p=1.233e-4
p=4.55e-1
p=4.50e-3
28
58
67
81
Tent5b-/- Tent5c-/- (n=48)
Tent5b wt/wt (n=66)
Tent5b wt/gfp (n=49)
Tent5b gfp/gfp (n=78)
Elobl
p=5.94e-07
p=1.00e-03
p=6.60e-10
21
53
66
75
Tent5b-/- Tent5c-/- (n=29)
Tent5b wt/wt (n=145)
Tent5b wt/gfp (n=131)
Tent5b gfp/gfp (n=214)
Zar1
p=4.71e-04
p=1.00e-03
p=7.53e-08
25
83
117
144
Tent5b-/- Tent5c-/- (n=27)
Tent5b wt/wt (n=91)
Tent5b wt/gfp (n=38)
Tent5b gfp/gfp (n=78)
Ezhip
p=2.460e-04
p=2.460e-04
p=1.842e-06
15
31
45
45
Tent5b-/- Tent5c-/- (n=24)
Tent5b wt/wt (n=76)
Tent5b wt/gfp (n=28)
Tent5b gfp/gfp (n=78)
Fbxw24
p=5.790e-09
p=1.190e-01
p=6.915e-04
31
128
150
187
Tent5b-/- Tent5c-/- (n=33)
Tent5b wt/wt (n=57)
Tent5b wt/gfp (n=45)
Tent5b gfp/gfp (n=39)
Nlrp14
p=1.008e-16
p=3.350e-01
p=1.500e-03
24
103
126
141
Tent5b-/- Tent5c-/- (n=79)
Tent5b wt/wt (n=123)
Tent5b wt/gfp (n=46)
Tent5b gfp/gfp (n=82)
Uchl1
p=1.854e-08
p=5.910e-04
p=8.040e-05
62
105
138
143
Tent5b-/- Tent5c-/- (n=123)
Tent5b wt/wt (n=219)
Tent5b wt/gfp (n=119)
Tent5b gfp/gfp (n=135)

**Supplemental Figure 3. TENT5s polyadenylate mRNAs in oocytes, which tight regulation is essential for oogenesis.**

DRS-based poly(A) tail lengths profiling of mRNAs isolated from ovaries. Median poly(A) tail lengths are plotted in white rectangles, p-values reported for comparison in Mann-Whitney-Wilcoxon test with Bonferroni Hallberg correction.

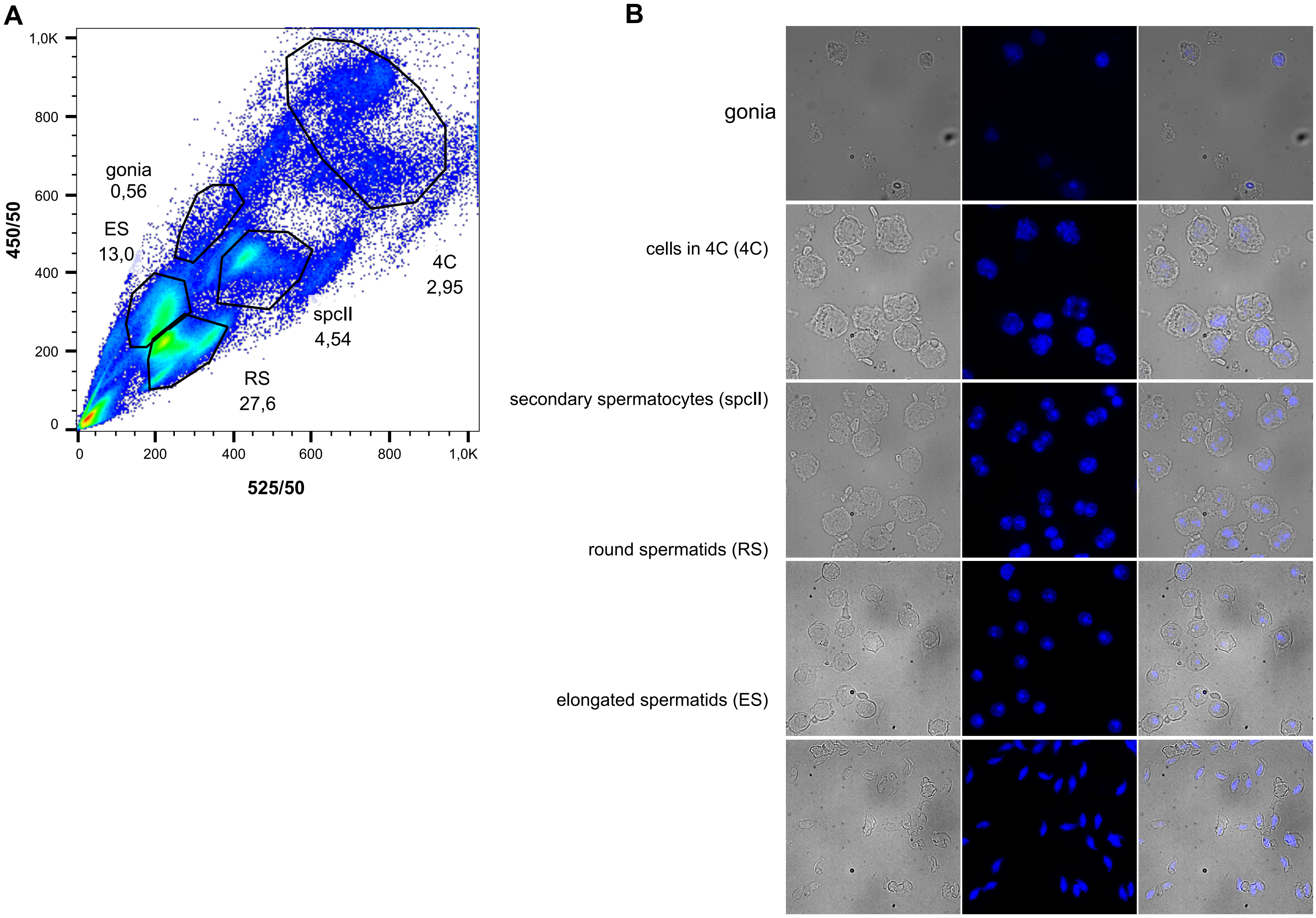


**Supplemental Figure 4. Male germ cells sorting**

**A**. Gating example of cells on different stages of spermatogenesis based on cells' size, shape, and DNA content in FACS analysis. **B**. Morphology and DNA staining pictures confirming cell sorting procedures correctness. Cells at different spermatogenesis stages display different patterns of DNA staining, size and shape.

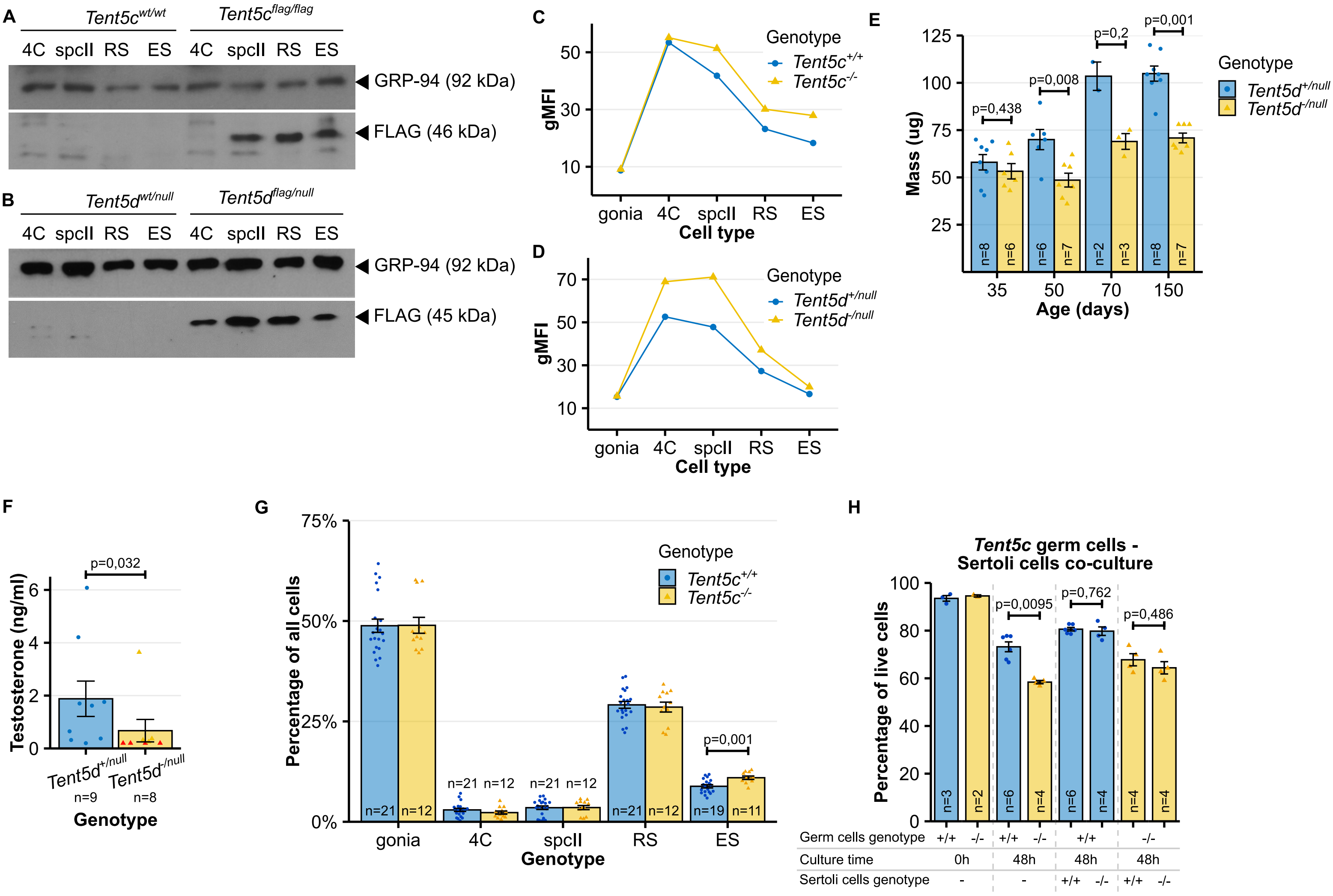
A
Tent5c wt/wt
Tent5c flag/flag
4C spcII RS ES 4C spcII RS ES
GRP-94 (92 kDa)
FLAG (46 kDa)
B
Tent5d wt/null
Tent5d flag/null
4C spcII RS ES 4C spcII RS ES
GRP-94 (92 kDa)
FLAG (45 kDa)
C
gMFI
Cell type
gonia 4C spcII RS ES
Genotype
Tent5c+/+
Tent5c-/-
D
gMFI
Cell type
gonia 4C spcII RS ES
Genotype
Tent5d+/null
Tent5d-/null
E
Mass (ug)
Age (days)
35 50 70 150
p=0,438
p=0,008
p=0,2
p=0,001
n=8 n=6 n=6 n=7 n=2 n=3 n=8 n=7
Genotype
Tent5d+/null
Tent5d-/null
F
Testosterone (ng/ml)
p=0,032
Tent5d+/null
Tent5d-/null
n=9
n=8
Genotype
G
Percentage of all cells
Genotype
Tent5c+/+
Tent5c-/-
n=21 n=12 n=21 n=12 n=21 n=12 n=21 n=12 n=19 n=11
p=0,001
gonia 4C spcII RS ES
Genotype
H
Tent5c germ cells -
Sertoli cells co-culture
Percentage of live cells
p=0,0095
p=0,762
p=0,486
n=3 n=2 n=6 n=4 n=6 n=4 n=4 n=4
Germ cells genotype +/+ -/- +/+ -/- +/+ -/-
Culture time 0h 48h 48h 48h
Sertoli cells genotype - - +/+ -/- +/+ -/-

**Supplemental Figure 5. Analysis of *Tent5c* and *Tent5d* KO phenotype in males**

**A-B.** Western blot analysis of FLAG-tagged TENT5C and TENT5D expression in different stages of spermatogenesis. Antibodies against GRP-94 were used as a loading control. Black arrowheads mark position of detected proteins. **C-D.** Cytometric analysis of GFP-tagged TENT5C and TENT5D expression in different stages of spermatogenesis, presented as changes as gMFI (geometric mean fluorescence intensity). **E.** Testes mass changes in $Tent5d^{+/null}$ and $Tent5d^{-/null}$ males in life. Individual data points and n values represent testes from individual males weighted, bars represent mean values, error bars represent SEM, p-values reported for comparison in Mann-Whitney-Wilcoxon test. **F.** Blood testosterone level in adult $Tent5d^{+/null}$ and $Tent5d^{-/null}$ males. Individual data points and n values represent samples from individual males, bars represent mean values, error bars represent SEM, p-values reported for comparison in Mann-Whitney-Wilcoxon test. **G.** Percentage of germ cells at different spermatogenesis stages among all isolated germ cells in $Tent5c^{+/+}$ and $Tent5c^{-/-}$ males. Individual data points and n values represent individual males from which germ cells were isolated, bars represent mean values, error bars represent SEM, p-value reported for comparison in t-test, two-tailed. **H.** Germ cell survival in 48h of *in vitro* culture depending on *Tent5c* genotype of germ cells and presence and genotype of Sertoli cells. Individual data points and n values represent cell culture of germ cells isolated from single male, bars represent mean values, error bars represent SEM; p-values reported for comparison in Mann-Whitney-Wilcoxon test.

**A**

**Motifs found in sequences of genes encoding potential TENT5D targets**

| region | motif | E-value | procentage of sequences maching the motive |
|---|---|---|---|
| 3'UTR | AAAGCTCACATATC | NN | 14.9% |
| | ATCACATCAAGCTCT | | 14.9% |
| | CCTCAAAGTCACACC | | 14.9% |
| | AGAGTTTTAAGTTTG | | 14.9% |
| | GGGGAAAAGA | | 23.4% |
| CDS | GTCTACCTATGATG | 3.0e+000 | 5.8% |
| | CCTGGCGCTT | 3.0e+000 | 11.5% |
| | ATTGCAGCGCCTTCA | 3.0e+000 | 5.8% |
| 5'UTR | CCCCGGCCTCCGGCC | NN | 27.8% |
| | ACCTCCGCCGCCACC | | 27.8% |
| | ACCCAGGCTGGA | | 27.8% |
| | AGGGTAGGGCGG | | 22.2% |
| | GCCCCCAGGCC | | 22.2% |

NN - Either the positive or negative hold-out set have been too small for STREME to reliably estimate E-value.

**B**

**Motifs found in sequences of genes encoding potential TENT5C targets**

| region | motif | E-value | procentage of sequences maching the motive |
|---|---|---|---|
| 3'UTR | CCAGGGGCG | 2.3e-003 | 9.4% |
| | TCCGGCACG | 5.9e-003 | 11.8% |
| | GGCCACCAGCG | 9.2e-002 | 10.6% |
| | ACTGTCCCT | 3.4e-001 | 10.6% |
| | GCGCGCAG | 6.6e-001 | 35.3% |
| CDS | CCCCAGGA | 8.1e-004 | 51.8% |
| | CCCCCGCCA | 1.3e-002 | 29.4% |
| | AGCAAAGAGATGT | 2.5e-002 | 3.2% |
| | CTGTGGGGCTGGC | 2.5e-002 | 2.5% |
| | CGAGCCAATAGTGGA | 5.1e-002 | 2.1% |
| 5'UTR | TCAGAATTCAGAAA | 1.9e-002 | 13.5% |
| | GCACCTCCCATGAC | 1.9e-002 | 6.8% |
| | GCCCGGGTAACGGCA | 3.8e-002 | 6.8% |
| | ATTCACCACC | 5.7e-002 | 5.4% |
| | ATTCACCACC | 7.0e+000 | 6.8% |

**C**

**Motifs found in sequences of genes encoding potential TENT5B and TENT5C**

| region | motif | E-value | procentage of sequences maching the motive |
|---|---|---|---|
| 3'UTR | AAGGCTGTG | 3.8e-002 | 18.2% |
| | ACATCCTCCAAA | 2.2e-001 | 18.2% |
| | CCGGGATC | 2.2e-001 | 15.3% |
| | AATAAAGGTCTATT | 1.2e+000 | 11.8% |
| | ACTAAAAGCA | 2.1e+000 | 12.9% |
| CDS | CAGTAAAGCAATC | 1.3e-002 | 4.2% |
| | TCCTCCAGAAAA | 2.5e-002 | 4.3% |
| | AGGAACAGAACTCAG | 2.6e-002 | 3.8% |
| | CACCCTACCTTTCC | 4.2e-002 | 1.6% |
| | GCCCTGTTCCT | 5.3e-002 | 6.8% |
| 5'UTR | ACACTGACTCTTC | 9.9e-002 | 9.9% |
| | ATGCCAGGCT | 1.4e-001 | 16.6% |
| | ATCCTAAGT | 3.0e-001 | 5.8% |
| | GGTCAGGAA | 3.0e-001 | 7.2% |
| | GAGAAGCCAA | 3.0e-001 | 8.5% |

**D**

**Biological procesess**

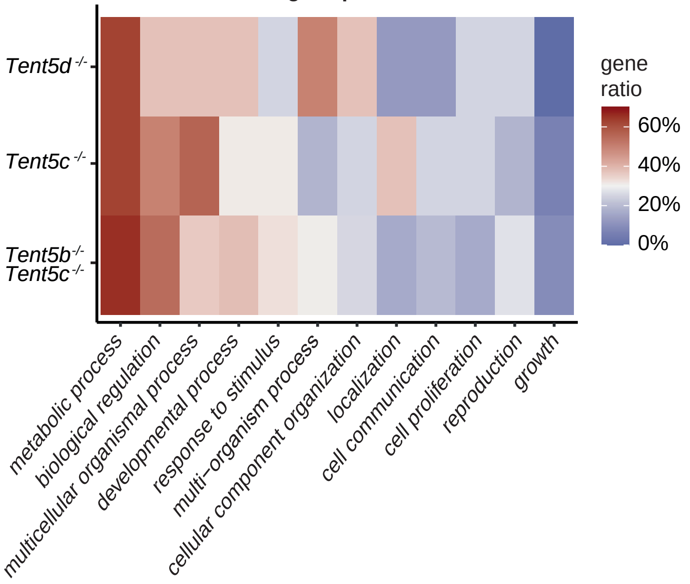

**Supplemental Figure 6. Motif and Gene ontology analysis of DRS results**

**A-C.** 3`UTR, 5’UTR motif analysis did not reveal any enriched motifs. In CDS only in Tent5C substrates, short C-rich motive is significantly enriched. **D-E.** Gene ontology (GO) analysis of potential TENT5 protein targets.

### Supplemental Table 1. Oligonucleotides, sgRNA and DNA repair templates sequences

| | |
|---|---|
| Fw primer for genotyping of Tent5a KO mouse line (#2001): CAAGCCTGATTGTGAAGGTG | Gewartowska et al.[29] |
| Rv primer for genotyping of *Tent5a* KO mouse line (#2002): AAGGAAGAGAAGGAAACGCA | Gewartowska et al.[29] |
| Fw primer for genotyping of *Tent5b* KO mouse line (large deletion) (#2007): TTAGCCTGAAGACGCTATGG | |
| Rv primer for genotyping of Tent5b KO mouse line (large deletion) (#2008): GCATGGGGGTCACATAGTAA | |
| Fw primer for genotyping of *Tent5b* KO mouse line (small deletion) (#2005): AATCGAGGCTTAGCGAGTTA | |
| Rv primer for genotyping of *Tent5b* KO mouse line (small deletion) (#2006) | |
| Fw primer for genotyping of *Tent5c* KO mouse line (#2012): AGGTCCTGACTGAGGTCGTG | Mroczek et al.[27] |
| Rv primer for genotyping of *Tent5c* KO mouse line (#2013): TTCCTCAAAATCCCCGTACA | Mroczek et al.[27] |
| Fw primer for genotyping of Tent5d KO mouse line (#2016): TCGGAGATCAGATTCAGCAAT | This paper |
| Rv primer for genotyping of Tent5d KO mouse line (#2017): TAGTTCTACGTTCTTCCCGTT | This paper |
| Fw primer for genotyping of *Tent5b*-GFP and *Tent5b*-FLAG mouse lines (#2310): GTGGTCAACGAGAGCACAGTG | This paper |
| Rv1 primer for genotyping of *Tent5b*-GFP and *Tent5b*-FLAG mouse lines (#2311): gaagatggtgcgctcctgg | This paper |
| Fw primer for genotyping of GFP-*Tent5b* mouse line (#2160): atcagaaactcctgagagcc | This paper |
| Rv1 primer for genotyping of GFP-*Tent5b* mouse line (#2180): GAATGGGGATCGGCTCTTTC | This paper |
| Rv2 primer for genotyping of GFP-*Tent5b* mouse line (#2326): gacacgctgaacttgtggc | This paper |
| Fw primer for genotyping of *Tent5c*-GFP and *Tent5c*-FLAG mouse lines (#2014): CTTCAGAACCACTTCTCGGA | Mroczek et al.[27] |
| Rv primer for genotyping of *Tent5c*-GFP and *Tent5c*-FLAG mouse lines (#2015): AGAAGTCACGCCTCCTATTG | Mroczek et al.[27] |
| Fw primer for genotyping of *Tent5d*-GFP and *Tent5d*-FLAG mouse lines (#2018): CCCAGTACAGACAACGTAAC | This paper |
| Rv primer for genotyping of *Tent5d*-GFP and *Tent5d*-FLAG mouse lines (#2019): GTGTTCTTTCATACGTTAGCC | This paper |
| gRNA used for generation of *Tent5b* KO mouse line: TGTAGCCTAGGCCGCTCTC | This paper |
| gRNA used for generation of *Tent5d* KO mouse line: ACATACTCGCAAGCCATAA | This paper |
| gRNA used for generation of *Tent5b*-GFP and *Tent5b*-FLAG mouse lines: AGGATCAGAGTCAGTTGCA | This paper |
| dsDNA used as repair template for generation of *Tent5b*-FLAG mouse line: ACCCCCATGCAGCCACTGCTGCCCCGAGCTCACTCCTATCCTACCTGGCTG CCTTGCAACGACTACAAAGACGATGACGACAAGTGACTCTGATCCTGGCCA GAAGGGAATGAGCGCCATGGGGTGGGGTGGGGTCATCAGGTA | This paper |

| | |
|---|---|
| dsDNA used as repair template for generation of *Tent5b*-GFP mouse line:<br>ACCCCCATGCAGCCACTGCTGCCCCGAGCTCACTCCTATCCTACCTGGCTG<br>CCTTGCAACgagaatttgtattttcagggtgatatcatggtgagcaagggcgaggagctgttcaccggggt<br>ggtgcccatcctggtcgagctggacggcgacgtaaacggccacaagttcagcgtgtccggcgagggcgagg<br>gcgatgccacctacggcaagctgaccctgaagttcatctgcaccaccggcaagctgcccgtgccctggccca<br>ccctcgtgaccaccctgacctacggcgtgcagtgcttcagccgctaccccgaccacatgaagcagcacgactt<br>cttcaagtccgccatgcccgaaggctacgtccaggagcgcaccatcttcttcaaggacgacggcaactacaa<br>gacccgcgccgaggtgaagttcgagggcgacaccctggtgaaccgcatcgagctgaagggcatcgacttca<br>aggaggacggcaacatcctggggcacaagCtggagtacaactacaacagccacaacgtctatatcatggc<br>cgacaagcagaagaacggcatcaaggtgaacttcaagatccgccacaacatcgaggacggcagcgtgca<br>gctcgccgaccactaccagcagaacacccccatcggcgacggccccgtgctgctgcccgacaaccactac<br>ctgagcacccagtccgccctgagcaaagaccccaacgagaagcgcgatcacatggtcctgctggagttcgt<br>gaccgccgccgggatcactctcggcatggacgagctgtacaagTGACTCTGATCCTGGCCAGA<br>AGGGAATGAGCGCCATGGGGTGGGGTGGGGTCATCAGGTA | This paper |
| dsDNA used as repair template for generation of GFP-*Tent5b* mouse line:<br>TTGGCCCGTGCACAGCCACTCTCCCTGCCCTCCTCGCCTTCACCATTCCC<br>GGGTTTCTGCCGTCCAGGCACCGGGGCCGGGTGAATGGTttCtAAGGGaGA<br>AGagctgttcacAggAgtggtgccTatcctggtcgagctggacggcgacgtaaacggccacaagttcagc<br>gtgtccggcgagggcgagggcgatgccacctacggcaagctgaccctgaagttcatctgcaccaccggcaa<br>gctgcccgtgccctggcccaccctcgtgaccaccctgacctacggcgtgcagtgcttcagccgctaccccgac<br>cacatgaagcagcacgacttcttcaagtccgccatgcccgaaggctacgtccaggagcgcaccatcttcttca<br>aggacgacggcaactacaagacccgcgccgaggtgaagttcgagggcgacaccctggtgaaccgcatcg<br>agctgaagggcatcgacttcaaggaggacggcaacatcctggggcacaagctggagtacaactacaacag<br>ccacaacgtctatatcatggccgacaagcagaagaacggcatcaaggtgaacttcaagatccgccacaaca<br>tcgaggacggcagcgtgcagctcgccgaccactaccagcagaacacccccatcggcgacggccccgtgct<br>gctgcccgacaaccactacctgagcacccagtccaagctgagcaaagaccccaacgagaagcgcgatca<br>catggtcctgctggagttcgtgaccgccgccgggatcactctcggcatggacgagctgtacaagggatCTgg<br>AGAAAACCTGTACTTCCAAGGAatgccATctgagagtggagctgaAagcctggagcagccag<br>ctgcgcaggtggggaccggtgcagcctcggcagtGGCCACGGCTG | This paper |
| dsDNA used as repair template for generation of *Tent5d*-FLAG mouse line:<br>CCACTGGTTTATTTCCAGCCATGTCATACAGTGCAGTTCCCTGTGCAAAATG<br>GTATGATGGACTACAAAGACGATGACGACAAGTAAGAAATACACATACCACA<br>AGTTTTGCTTAAGCAACTCTGAAAAAGCAATTTTCCAAGT | This paper |
| dsDNA used as repair template for generation of *Tent5d*-GFP mouse line:<br>CCACTGGTTTATTTCCAGCCATGTCATACAGTGCAGTTCCCTGTGCAAAATG<br>GTATGATGgagaatttgtattttcagggtgatatcatggtgagcaagggcgaggagctgttcaccggggtg<br>gtgcccatcctggtcgagctggacggcgacgtaaacggccacaagttcagcgtgtccggcgagggcgaggg<br>cgatgccacctacggcaagctgaccctgaagttcatctgcaccaccggcaagctgcccgtgccctggcccac<br>cctcgtgaccaccctgacctacggcgtgcagtgcttcagccgctaccccgaccacatgaagcagcacgacttc<br>ttcaagtccgccatgcccgaaggctacgtccaggagcgcaccatcttcttcaaggacgacggcaactacaag<br>acccgcgccgaggtgaagttcgagggcgacaccctggtgaaccgcatcgagctgaagggcatcgacttcaa<br>ggaggacggcaacatcctggggcacaagctggagtacaactacaacagccacaacgtctatatcatggccg<br>acaagcagaagaacggcatcaaggtgaacttcaagatccgccacaacatcgaggacggcagcgtgcagc<br>tcgccgaccactaccagcagaacacccccatcggcgacggccccgtgctgctgcccgacaaccactacctg<br>agcacccagtccgccctgagcaaagaccccaacgagaagcgcgatcacatggtcctgctggagttcgtgac<br>cgccgccgggatcactctcggcatggacgagctgtacaagTAAGAAATACACATACCACAAGT<br>TTTGCTTAAGCAACTCTGAAAAAGCAATTTTCCAAGT | This paper |
| Tn5ME-A:<br>CGTCGGCAGCGTCAGATGTGTATAAGAGACAG | Hennig et al.[64] |
| Tn5ME-B:<br>GTCTCGTGGGCTCGGAGATGTGTATAAGAGACAG | Hennig et al.[64] |
| Tn5MErev:<br>[phos]CTGTCTCTTATACACATCT | Hennig et al.[64] |
| Fw primer for SLIC cloning:<br>atgaacgagctctataagagatctttcgaaaaggacgagctgtaacaaagaaagcccagctccttc | This paper |
| Rv primer for SLIC cloning:<br>cacagtcgaggctgatcagcgggtttaaacaaagtgtgaaaaatacctctg | This paper |

| | |
|---|---|
| Fw oligonucleotide for Zp3 signal peptide sequence: agcttATGGCGTCAAGCTATTTCCTCTTCCTTTGTCTCCTGCTGTGTGGAGGCCCCGAGCTGTGCAATTCC | This paper |
| Rv oligonucleotide for Zp3 signal peptide sequence: CCGGGGAATTGCACAGCTCGGGGCCTCCACACAGCAGGAGACAAAGGAAGAGGAAATAGCTTGACGCCATa | This paper |
| Fw oligonucleotide for Gdf9 signal peptide sequence: agcttATGGCACTTCCCAGCAACTTCCTGTTGGGGGTTTGCTGCTTTGCCTGGCTGTGTTTTCTTAGTAGCCTTAGCTCTCAGGCTTCTACT | This paper |
| Rv oligonucleotide for Gdf9 signal peptide sequence: CCGGAGTAGAAGCCTGAGAGCTAAGGCTACTAAGAAAACACAGCCAGGCAAAGCAGCAAACCCCCAACAGGAAGTTGCTGGGAAGTGCCATa | This paper |
| Fw primer for plasmid linearization: tagagaacccactgcttactgg | This paper |
| Rv primer for plasmid linearization: TTTTTTTTTTTTTTTTTTTTggattgaaggagctgggcttt | This paper |
